# Genetic architecture of a body color cline in *Drosophila americana*

**DOI:** 10.1101/2020.05.07.074211

**Authors:** Lisa L. Sramkoski, Wesley N. McLaughlin, Arielle M. Cooley, David C. Yuan, Alisha John, Patricia J. Wittkopp

## Abstract

Phenotypic variation within a species is often structured geographically in clines. In *Drosophila americana*, a longitudinal cline for body color exists within North America that appears to be due to local adaptation. The *tan* and *ebony* genes have been hypothesized to contribute to this cline, with alleles of both genes that lighten body color found in *D. americana*. These alleles are similar in sequence and function to the allele fixed in *D. americana’s* more lightly pigmented sister species, *Drosophila novamexicana*. To test this hypothesis, we examined the frequency and geographic distribution of *D. novamexicana*-like alleles of *tan* and *ebony* in *D. americana*. Among alleles from over 100 strains of *D. americana* isolated from 21 geographic locations, we failed to identify additional alleles of *tan* or *ebony* with as much sequence similarity to *D. novamexicana* as the alleles previously described. However, using genetic analysis of 51 *D. americana* strains derived from 20 geographic locations, we identified one new allele of *ebony* and one new allele of *tan* segregating in *D. americana* that are functionally equivalent to the *D. novamexicana* allele. An additional 5 alleles of *tan* also showed marginal evidence of functional similarity. Given the rarity of these alleles, however, we conclude that they are unlikely to be driving the pigmentation cline. Indeed, phenotypic distributions of the 51 backcross populations analyzed indicate a more complex genetic architecture, with diversity in the number and effects of loci altering pigmentation observed both within and among populations of *D. americana*. This genetic heterogeneity poses a challenge to association studies and genomic scans for clinal variation, but might be common in natural populations.

## Introduction

A phenotypic cline describes a gradient of trait variation across geographic space (Huxley 1938). Such clinal variation often correlates with latitude, longitude or altitude, which in turn correlate with environmental factors such as temperature, light, and humidity. Clinal trait variation can arise neutrally from reduced gene flow between geographically distant populations, but natural selection favoring adaptation to varying local environments is more often thought to be responsible -- especially when there is ongoing gene flow among populations (Endler 1977). Genetic variation underlying clinal trait variation is frequently sought by searching for matching allele frequency clines, but this strategy is known to produce many false positives (Lotterhos & Whitlock 2015; François *et al.* 2016). Incorporating knowledge of gene function can help overcome this limitation by identifying loci most likely to contribute to trait variation (Stinchcombe & Hoekstra 2007; Fournier-Level *et al.* 2011; Hancock *et al.* 2011; Marjoram *et al.* 2013). Genome scans can also miss loci contributing to clinal trait variation when traits are controlled by many genes: for such polygenic traits, multiple genotypes can often produce the same phenotype (genetic heterogeneity), which complicates expected allelic variation across a cline (Kawecki & Ebert 2004; Pritchard & Di Rienzo 2010; Savolainen *et al.* 2013; Adrion *et al.* 2015; Haasl & Payseur 2016). Here, we use a more targeted approach to investigate the genetic basis of clinal trait variation by directly examining the role of two genes known to affect development of a clinally varying, polygenic trait. More specifically, we examine the contributions of divergent *tan* and *ebony* alleles to clinal variation of body color in *Drosophila americana*.

The genetic basis of pigmentation differences within and between species has been studied extensively within Drosophila (Massey & Wittkopp 2016), and pigmentation clines for body color have been reported for many species (e.g., David *et al.* 1985; David & Capy 1988; Hollocher *et al.* 2000; Pool & Aquadro 2007; Wittkopp *et al.* 2011; Telonis-Scott *et al.* 2011). Selection pressures driving these pigmentation clines seem to vary among species, with adaptation proposed to be linked to variation in UV radiation, temperature, and/or humidity (David & Capy 1988; True 2003; Brisson *et al.* 2005; Rajpurohit *et al.* 2008; Wittkopp & Beldade 2009; Clusella-Trullas & Terblanche 2011; Parkash *et al.* 2012; Matute & Harris 2013; Bastide *et al.* 2014; Sillero *et al.* 2014; Rajpurohit & Schmidt 2019; Davis & Moyle 2019). In *D. americana*, which is found in the United States from the Atlantic coast to just east of the Rocky Mountains, pigmentation varies along a longitudinal cline, with the darkest body color seen among the most eastern populations (Wittkopp *et al.* 2011). This pigmentation cline is observed despite little evidence of population structure in *D. americana* and signatures of extensive gene flow throughout the species range (Schäfer *et al.* 2006; Morales-Hojas *et al.* 2008; Fonseca *et al.* 2013), suggesting it is due to local adaptation (Wittkopp *et al.* 2011). *D. americana*’s closest living relative, *D. novamexicana*, is found in the southwestern United States, west of the Rocky Mountains, and has evolved an even lighter body color, consistent with an extension of the *D. americana* pigmentation cline (Wittkopp *et al.* 2011). Although *D. americana* and *D. novamexicana* show evidence of reproductive isolation (Ahmed-Braimah & McAllister 2012), these two species are still able to mate and produce fertile offspring in the lab, allowing genetic dissection of their divergent phenotypes.

Pigmentation differences between *D. americana* and *D. novamexicana* have been linked to divergent alleles of two classic pigmentation genes, *ebony* and *tan,* with genomic regions containing these two genes explaining ~87% of the pigmentation difference (Wittkopp *et al.* 2003; 2009; Cooley *et al.* 2012). Proteins encoded by *ebony* and *tan* are required for pigment synthesis in Drosophila and catalyze opposite directions of a reversible biochemical reaction converting dopamine to N-beta-alanyl dopamine and vice versa (True *et al.* 2005; Massey & Wittkopp 2016). For *tan*, functionally divergent sites have been mapped to the first intron (Wittkopp *et al.* 2009) and allele-specific expression analysis in F_1_ hybrids (Wittkopp *et al.* 2004) suggests that this divergence affects *cis*-regulation of *tan* expression (Cooley *et al.* 2012). Evidence of *cis*-regulatory divergence between *D. americana* and *D. novamexicana* has also been detected for *ebony* using allele-specific expression assays (Cooley *et al.* 2012); however, the specific sites responsible for this divergence have been difficult to localize because *ebony* is located in a region of the genome inverted between *D. novamexicana* and *D. americana* (Wittkopp *et al.* 2009). Recent work using CRISPR/Cas9 genome editing to generate *ebony* mutants in both *D. americana* and *D. novamexicana,* however, has shown using reciprocal hemizygosity testing that divergent *ebony* alleles are indeed responsible for pigmentation differences between these two species (Lamb *et al.* 2020).

The contribution of *ebony* and *tan* to pigmentation differences between *D. americana* and *D. novamexicana* suggests that one or both of these genes might also contribute to variable pigmentation within *D. americana*. Consistent with this possibility, prior work identified a strain of *D. americana* (DN2) with an allele of *ebony* that shares both sequence and function with the *D. novamexicana* allele (Wittkopp *et al.* 2009). A different strain of *D. americana* (A01) was found to carry an allele of *tan* with sequence and function similar to the *D. novamexicana* allele (Wittkopp *et al.* 2009). These alleles seem to have arisen prior to speciation (Wittkopp *et al.* 2009), suggesting that they were segregating in *D. americana* prior to the divergence of *D. novamexicana*. Based on these data, we hypothesized that differences in the frequency of one or both of these *D. novamexicana*-like alleles among *D. americana* populations might contribute to this species’ pigmentation cline. Here, we test this hypothesis by searching over 100 strains of *D. americana* for additional alleles of *ebony* and/or *tan* that share similar amounts of sequence identity and/or function to the *D. novamexicana* allele. We then test for associations between pigmentation and segregating sites sampled in *ebony* and *tan*. Finally, we analyze pigmentation phenotypes of backcross populations between *D. novamexicana* and 51 strains of *D. americana* to determine how the genetic architecture of body color differs among strains. We find that *D. novamexicana*-like alleles of *ebony* and *tan* are unlikely to explain the body color cline in *D. americana,* and that the genetic architecture is more complex than anticipated, with genetic heterogeneity apparently common within populations affected by local adaptation. These observations suggest that genomic scans for variation in allele-frequencies would fail to find loci underlying this phenotypic cline, as has been predicted for clinally varying polygenic traits (Pritchard & Di Rienzo 2010; Savolainen *et al.* 2013; Adrion *et al.* 2015; Haasl & Payseur 2016).

## Materials and Methods

### Fly strains used for sequence analysis

A summary of fly strains used for sequence analysis is provided in Supplementary Table 1. The “A01” strain of *D. americana* (15010-0951.01) and “N14” strain of *D. novamexicana* (15010-1031.14) were obtained from the Drosophila Species Stock Center (Tucson, AZ). The remaining 112 strains of *D. americana* were generously provided by Dr. Bryant McAllister (University of Iowa), who collected the progenitors of these isofemale lines from wild populations between 1996 and 2007 at 21 sites sampled within the population range of *D. americana* in the United States. All flies were reared on a diet of standard yeast-glucose media at 20°C. Please note that we refer to different collection sites as different populations in the main text for simplicity even though patterns of sequence variation show no evidence of population structure in *D. americana* other than for chromosomal fusions and inversions (Schäfer *et al.* 2006; Morales-Hojas *et al.* 2008; Wittkopp *et al.* 2011; Fonseca *et al.* 2013).

### DNA sequence analysis

We PCR amplified and Sanger sequenced 579 bp of *ebony* spanning exons 5-8 and 1328 bp of *tan* from intron 1. (Note that we originally targeted the large first intron of *ebony*, but polymorphisms among strains caused all primer pairs tested to amplify inconsistently among strains.) After removing low quality bases from raw Sanger sequence reads based on Phred scores, we aligned sequences of *ebony* from 109 strains of *D. americana* plus 1 strain of *D. novamexicana* and sequences of *tan* from 102 strains of D. americana plus 1 strain of *D. novamexicana* using the ClustalW algorithm (Thompson *et al.* 1994) in CodonCode Aligner (version 8.0.2, https://www.codoncode.com/); sequence was obtained for both genes from 99 strains of *D. americana* (Supplementary Table 1). Only a single strain of *D. novamexicana* was analyzed in this work because prior work has shown very low levels of polymorphism in this species (Orsini *et al.* 2004; Caletka & McAllister 2004; Wittkopp *et al.* 2009). Sequence alignments used for analysis are provided as Supplementary File 1(*ebony*) and Supplementary File 2 (*tan*) and were submitted to GenBank with ID numbers MT350927 - MT351036 for *ebony* and MT350824 - MT350926 for *tan*.

### Gene trees and haplotype network analysis

Phylogenetic trees inferring evolutionary relationships among the alleles sampled for *ebony* and *tan* were produced using the Maximum Likelihood method based on the Tamura-Nei model of nucleotide substitutions (Tamura & Nei 1993) in MEGA7 (Kumar *et al.* 2016). A bootstrap consensus tree was inferred from 100 replicates (Felsenstein 1985), with branches supported by less than 50% of the replicates collapsed. As described in MEGA7, trees used to start the heuristic search were generated using the Neighbor-Join and BioNJ algorithms, with pairwise distances estimated using the Maximum Composite Likelihood (MCL) approach. Topologies with superior log likelihood values were then selected as initial trees. Sites for which 5% of the strains had alignment gaps, missing data, or ambiguous bases were excluded from this analysis. Because linkage disequilibrium is low within *D. americana* (Wittkopp *et al.* 2009), we also assessed the sequence similarity among alleles using Median Spanning Networks (Bandelt *et al.* 1999) (as implemented in PopART (www.popart.otago.ac.nz; March 15, 2015 version, downloaded September 12, 2019) with the epsilon parameter set to 0.

### Fly strains used for genetic analysis

The genetic basis of pigmentation differences between *D. americana* and *D. novamexicana* was examined for 51 of the *D. americana* strains established and provided by Dr. Bryant McAllister (University of Iowa) (McAllister *et al.* 2008; Sheeley & McAllister 2008). As shown in Supplementary Table 1, these strains of *D. americana* included 5 strains from each of two locations, 4 strains from each of two locations, 3 strains from each of six locations, 2 strains from each of five locations, and 1 strain from each of five locations. The eastern-most location was Killbuck, Ohio (40.711809, −82.005472), the western- and northern-most location was Niobrara, Nebraska (42.74821, −98.051519), and the southern-most collection site was Sneads, Florida (30.708495, −84.910637). Together, these 51 strains came from 20 of the 21 locations from which strains included in the sequence analysis described above were derived (Supplementary Table 1).

### Fly crosses for genetic analysis

Virgin females were isolated from each of the 51 strains of *D. americana* used for genetic analysis and mated with *D. novamexicana* males to create F_1_ hybrids. From each of these F_1_ hybrid populations, virgin females were again collected and then mated to *D. novamexicana* males. Male flies were collected from the (BC_1_) progeny produced by each backcross within 3 days of eclosion and aged for one week to ensure pigmentation was fully developed. Each of these BC_1_ males carried an X chromosome and one copy of each autosome that was a unique recombination of alleles from the *D. novamexicana* and *D. americana* strains crossed to generate its F_1_ hybrid mother. These different recombinant chromosomes caused pigmentation to vary among BC_1_ flies from each cross. The Y chromosome and the other copy of each autosome in the BC_1_ males was always inherited from the *D. novamexicana* father.

### Phenotyping Pigmentation in Backcross Progeny

For each backcross population, pigmentation of 27 to 117 (mean = 63.5) male BC_1_ flies 7-10 day old were scored based on the color visible in the dorsal abdominal cuticle of live flies. We found that pigmentation phenotypes did not vary continually in these backcross populations, but rather fell into distinct classes, consistent with prior work (Wittkopp *et al.* 2003; 2009). The number of distinct pigmentation classes used to score each backcross population was based on the number of distinct pigmentation phenotypes observed: we observed four to eight distinct classes of pigmentation phenotypes in each of the 51 BC_1_ populations. The lightest class was always designated as category “1” with increasing class numbers corresponding to progressively darkening pigmentation. For example, in a backcross population with four total pigmentation classes, class “4” would contain the darkest flies, whereas in a backcross population with seven total pigmentation classes, class “4” would contain flies with mid-range pigmentation. The number of pigmentation classes as well as the assignment of individual flies to a particular pigmentation class was determined by independent observations from at least two researchers. These pigmentation phenotype scores are shown for each fly in Supplementary Table 3.

### DNA Extractions

From each of the 51 backcross populations, DNA was extracted from each male BC_1_ fly using a method similar to that described in Gloor *et al.* (1993) except that the protocol was scaled for efficient processing of 3238 flies. Briefly, each fly was placed into a well of a 96-well plate (GeneMate# T3031-21) with a glass bead and 50μL of a 1:99 Proteinase K/Engel’s Buffer solution. Plates were sealed and shaken in a Qiagen Retsch MM301 Tissue Lyser until the glass bead had pulverized the fly in each well. The plates were then incubated at 37°C for 30 minutes to allow protein digestion and then incubated at 95°C for 2 minutes to inactivate Proteinase K. Extracted DNA was stored at 4°C until used for genotyping.

### Genotyping

Molecular genotyping assays were used to determine whether each of the BC_1_ males scored for pigmentation carried the *D. americana* and/or *D. novamexicana* alleles of three pigmentation genes: *yellow*, *tan*, and *ebony*. Because *yellow* and *tan* are located on the X chromosome, each male carried only one species’ allele, either the mother’s or the father’s allele. By contrast, because *ebony* is located on an autosome, BC_1_ males could either be heterozygous for the *D. americana* and *D. novamexicana* alleles or homozygous for the *D. novamexicana* allele.

For *yellow* and *tan*, differences in length between PCR products amplified from the *D. americana* and *D. novamexicana* alleles were used to genotype BC1 flies. For *tan,* a forward primer (5’-CGAGTTTTTATTCCCACTGAATTAT-3’) and a reverse primer (5’-GGGTTCGTCTTATCCACGAT-3’) were used to amplify a 100bp product for the *D. americana tan* allele and a 64bp product for the *D. novamexicana tan* allele. For *yellow,* depending on which *D. americana* strains was used to generate the BC_1_ males being genotyped, one of two forward primers was used [*yellow* forward-1 (5’-CCAAAAGGACAACCGAGTTT-3’) or *yellow* forward-2 (5’-CTAAACATGCCTGAAAATCAATCACGGA-3’)] with a *yellow* reverse primer (5’-AGTCGATTGCCAAAGTGCTC-3’). These different forward primers were necessary because of differences in *yellow* DNA sequence among the *D. americana* strains. For most backcross populations, the *yellow* forward-1 primer paired with the *yellow* reverse primer generated a 349bp product for the *D. americana yellow* allele and a 372bp product for the *D. novamexicana yellow* allele. The *yellow*-forward-2 primer was used to analyze BC_1_ males from the six strains of *D. americana* (IR0436, LR0540, FP9946, DI0562 MK0738, and SC0708) for which the *yellow* forward-1 primer and *yellow* reverse primer did not produce any visible differences in length between the *D. americana* and *D. novamexicana* alleles. For these six strains, genotyping was performed by using the *yellow* forward-2 primer and the *yellow* reverse primer to amplify a region of *yellow* using PCR and then digesting the PCR product with DraI, which cut only the *D. novamexicana yellow* allele. All digested and undigested PCR products were run on 2% agarose gels and visualized using Ethidium Bromide.

For *ebony*, we were unable to identify PCR products that were easily distinguishable for *D. americana* and *D. novamexicana* alleles through either amplicon length or restriction digest. Therefore, we genotyped flies at the *ebony* locus using pyrosequencing (Ahmadian *et al.* 2000). The PCR product used for pyrosequencing was generated using the forward primer, 5’-AGCCCGAGGTGGACATCA-3’, and the biotinylaed reverse primer, 5’-*GTATGGGTCCCTCGCAGAA-3’ (* notates biotinylation). These PCR products were processed, and pyrosequencing performed, as described in Wittkopp *et al.* (2008). The pyrosequencing primer used had the sequence 5’-CGAGGTGGACATCAAGT-3’. This pyrosequencing assay for *ebony* used two single nucleotide differences to differentiate between the *D. americana* and *D. novamexicana ebony* alleles. Specifically, the sequences analyzed by pyrosequencing were 5’-C**C**AAGCT**G**CT-3’ for the *D. americana* allele and 5’-C**G**AAGCT**T**CT-3’ for the *D. novamexicana* allele, where the bolded letters indicate bases used to discriminate between the two alleles.

Genotyping data for *yellow, tan*, and *ebon*y in the BC_1_ males is summarized in Supplementary Table 4, where 0 = hemizygous for the *D. americana* allele for *yellow* and *tan* and heterozygous for *ebony* and 1 = hemizygous *D. novamexicana* allele for *yellow* and *tan* and homozygous for *ebony.* The 96-well plate containing the DNA sample from each fly is also indicated in Supplementary Table 4.

### Comparing function of *D. americana ebony, tan*, and *yellow* alleles to *D. novamexicana*

To determine whether the *D. americana* allele of *yellow*, *tan*, and/or *ebony* from each of the 51 strains of *D. americana* examined was functionally equivalent to the *D. novamexicana* allele of the same gene, we calculated the difference between the mean pigmentation scores of flies inheriting the *D. americana* or *D. novamexicana* allele from their mother in each backcross population. Statistical significance of this difference was assessed for each gene in each backcross using a null distribution of pigmentation differences generated from 10,000 permuted datasets in which the genotypes of the focal gene were shuffled relative to the pigmentation phenotypes. The null hypothesis tested by these permutations was that the *D. americana* and *D. novamexicana* alleles of the focal gene had indistinguishable effects on pigmentation (i.e., that the two alleles are functionally equivalent). This method of testing for statistical significance directly accounts for the differences in sample sizes and allele frequencies among backcrosses. A correction for multiple testing was performed with the p.adjust function with the method=fdr option, which implements the false discovery rate correction as described in Benjamini & Hochberg (1995). These adjusted p-values are reported in Supplementary Table 5.

### Association testing

To test for an association between pigmentation and segregating sites in *tan* and *ebony*, we used a more quantitative, continuous measure of pigmentation than the pigmentation classes described for backcross populations above. This pigmentation data came from dataset B in Wittkopp et al. (2011) for strains from the DN, Il, MK, NN, OC, SC, and WS populations. For the remaining strains, we generated comparable quantitative measurements of pigmentation using the same protocol as described for dataset B in Wittkopp et al. (2011). Briefly, a custom-built fiber optic probe was used to measure light reflected off the fly’s abdominal cuticle, with 5 measurements taken per fly and 6-20 flies analyzed per strain. A WS-1 Diffuse Reflection Standard (Ocean Optics) was used to calibrate the probe for each set of measurements and strains were scored in a random order. To minimize the effects of outlier measurements, the median measure of pigmentation observed for each fly was used for analysis. These medians (Supplementary Table 2) were fitted to a linear model including strain as a fixed effect and replicate fly as a random effect with *lmer* function in the *lme4* R package, and the least-square means were extracted for each strain using the *lsmeans* function in the *lsmeans* R package.

Variable sites were then identified in *tan* and *ebony* using the same sequence alignments used for phylogenetic analysis (Supplementary Files 1 and 2). Sites with the minor allele present in less than 5 strains as well as sites containing indels were excluded prior to association testing. Each of the remaining variable sites for *tan* (N = 74) and *ebony* (N = 40) was then tested for an association with pigmentation by fitting the lsmean estimate of pigmentation for each strain to a general linear model (function *glm* in R) containing each of the variable sites as a fixed effect.

### Standardizing pigmentation classes among strains

One representative male fly from each phenotypic class in each backcross was imaged as a visual reference using a Scion Visicapture 1.2 and Scion Corporation Model CFW-1308C color digital camera. These images were processed using Photoshop CS6 (Adobe, San Jose, CA), with a constant color adjustment applied to all photos collected on the same day to control for day-to-day variation in imaging conditions. These adjustments were performed to make the digital images more closely match the fly’s appearance under the microscope. The parameters for each day’s adjustment were determined based on images of a set of standards consisting of seven dissected abdominal cuticles with a range of pigmentation phenotypes. Photos of these cuticle standards were collected interleaved within each batch of BC_1_ flies. For comparisons among flies from all 51 backcross populations, we used the representative images from each category in each backcross to convert backcross-specific pigmentation scores to a common 8-category pigmentation scale (Supplementary Table 3). After phenotyping, all flies were stored at −80°C.

### Comparing distributions of backcross phenotypes among strains

Correspondence analysis (CA), which is similar to principal components analysis but for categorical response variables, was used to reduce the dimensionality of the distributions of pigmentation classes from backcross (BC1) populations among strains. This analysis was performed using the *CA* function in the *FactoMineR* package (Lê *et al.*, 2008) for R and visualized using *factoextra* R package. We then calculated the Euclidean distance between strains in the Dimension 1 and Dimension 2 space from the CA analysis to compare the similarity in backcross pigmentation distributions for strains that were and were not from the same collection site. Euclidean distances between all pairs of strains were calculated using the *distances* function in the *distances* R package.

### Statistical analyses

R code used for this work is provided in Supplementary File 3. This code was run in RStudio (Version 1.2.5033) using R version 3.6.2 (2019-12-12).

## Results

### Comparing sequence of *D. americana ebony* and *tan* alleles to *D. novamexicana* alleles

As described in the Introduction, pigmentation differences between *D. americana* and *D. novamexicana* (Figure 1A) are primarily due to changes in the *ebony* and *tan* genes, which control the balance between dark (black and brown) and light (yellow/tan) pigments (Figure 1B). The DN2 strain of *D. americana* (from Duncan, Nebraska) and the A01 strain of *D. americana* (from Poplar, Montana) have been shown to carry alleles of *ebony* and *tan*, respectively, similar in sequence and function to the *D. novamexicana* alleles of these genes (Wittkopp *et al.* 2009). These observations suggest that differences in the frequency of *D. novamexicana*-like alleles among populations of *D. americana* might underlie the longitudinal cline of body color observed within this species. To test this hypothesis, we examined the frequency and geographic distribution of such alleles first by comparing sequences of *ebony* and *tan* from over 100 strains of *D. americana* to orthologous sequences from the N14 strain of *D. novamexicana*. The *D. americana* strains examined were derived from flies captured at 21 different sites within the United States and included DN2 and A01 (Figure 1C, Supplementary Table 1).

**Figure 1.**
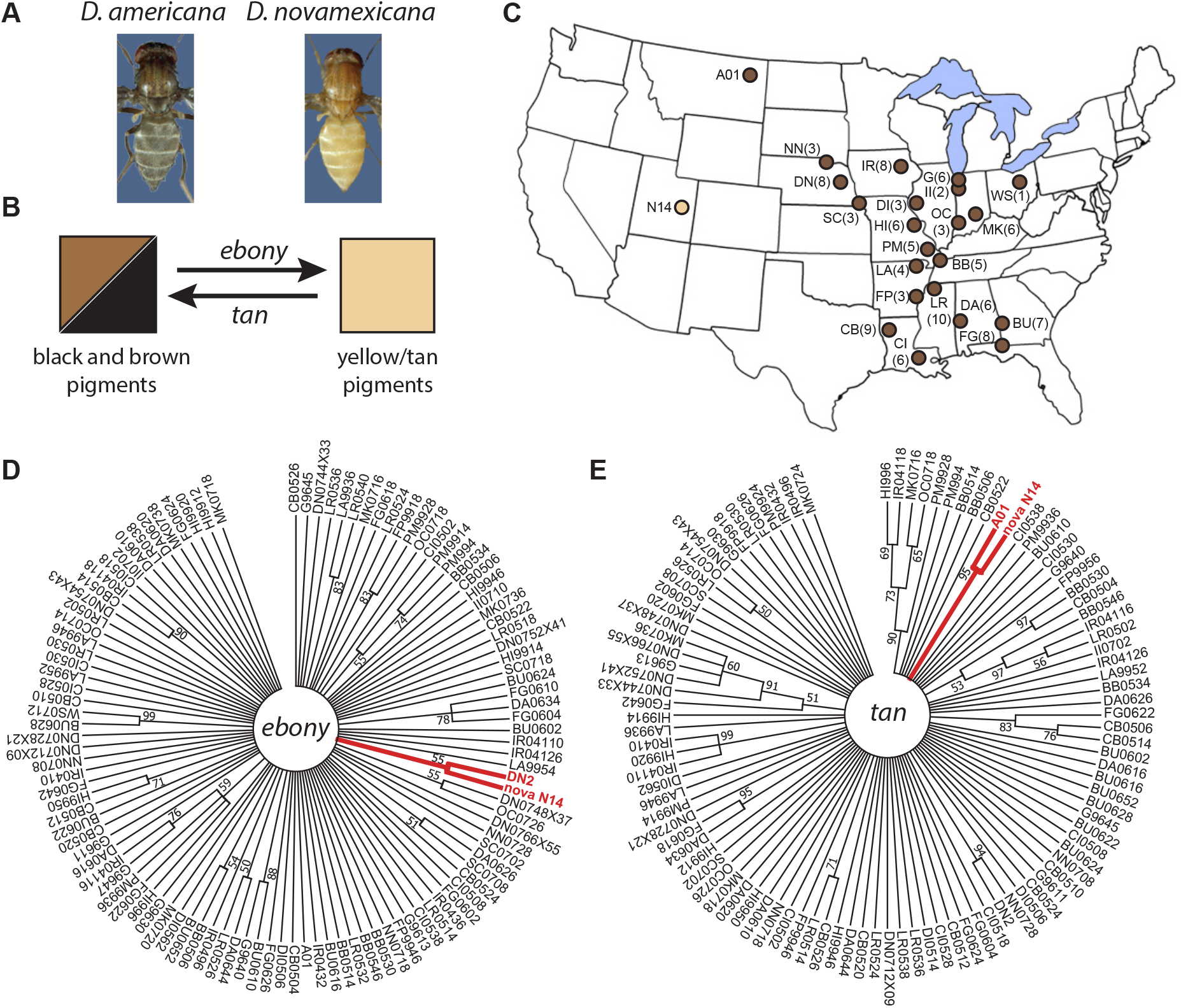
*D. americana* alleles of *ebony* and *tan* closely related to the *D. novamexicana* allele are rare within *D. americana.* (A) *D. americana* (left) has a much darker body color than *D. novamexicana* (right). (B) The *tan* and *ebony* genes encode enzymes that catalyze a reversible biochemical reaction required for the production of dark (black and brown) melanins and light (yellow/tan) sclerotins, respectively. (C) Collection sites for progenitors of *D. americana* (brown) and *D. novamexicana* (yellow) strains used in this work are shown. Numbers in parentheses indicate the number of independently isolated strains examined from that site. Only a single strain from the Drosophila Species Stock Center was examined for A01 and N14. For more information about these strains, see Supplementary Table 1. (D, E) The circular phylogenetic trees shown for *ebony* (D) and *tan* (E) were produced using a Maximum Likelihood method implemented in MEGA7, as described in Methods. Branches shown were supported by 50% or more of bootstrap replicate trees. The *ebony* tree is based on 579 aligned sites from 110 alleles, and the *tan* tree is based on 1328 aligned sites from 103 alleles. Branches shown in red highlight the *D. novamexicana* allele (“nova N14”) and the allele from *D. americana* (DN2 for *ebony*, A01 for *tan*) previously shown to share similarity in both sequence and function with the *D. novamexicana* allele (Wittkopp *et al.* 2009).

Phylogenetic trees built from these sequences using the maximum likelihood method implemented in MEGA7 (Kumar *et al.* 2016) confirmed that the *ebony* allele from the DN2 strain of *D. americana* is more similar to the *D. novamexicana* allele than to other alleles from *D. americana* (Figure 1D). We failed to find, however, any additional *ebony* alleles from the 109 new strains of *D. americana* sampled that clustered as closely with *D. novamexicana* (Figure 1D). Similarly, phylogenetic trees confirmed that the *tan* allele from the A01 strain of *D. americana* was the only allele among those sampled from 102 strains of *D. americana* that is more closely related to the *D. novamexicana* allele than to other *D. americana* alleles (Figure 1E). Analyzing these sequences with Minimum Spanning Networks implemented in PopArt (www.popart.otago.ac.nz) also showed that the DN2 and A01 alleles of *ebony* and *tan*, respectively, were most similar to the *D. novamexicana* allele (Supplementary Figures 1 and 2). Taken together, these data indicate that alleles of *ebony* and *tan* with sequences closely related to the *D. novamexicana* allele are rare within *D. americana* and thus unlikely to explain the pigmentation cline observed.

### Comparing function of *D. americana ebony* and *tan* alleles to *D. novamexicana* alleles

To determine whether other *D. americana* alleles of *ebony* and/or *tan* might have functional similarity to *D. novamexicana* alleles despite their greater sequence divergence, we crossed virgin females from 51 strains of *D. americana* derived from 20 populations (Supplementary Table 1) to *D. novamexicana*, and then backcrossed the F_1_ hybrid females to *D. novamexicana* males (Figure 2A). The backcross (BC1) progeny inherited recombinant maternal chromosomes that contain sequences from both their *D. americana* and *D. novamexicana* parents and paternal chromosomes with only *D. novamexicana* alleles (Figure 2A). Pigmentation was scored for all male flies in each backcross population (N = 27 to 117, mean = 63.5), and then each male was genotyped for *ebony*, *tan*, and another pigmentation gene, *yellow* (Supplementary Table 2). The *yellow* gene was included as a negative control in this study because prior work has shown that it does not contribute to pigmentation divergence between *D. americana* and *D. novamexicana* (Wittkopp *et al.* 2003; 2009).

**Figure 2.**
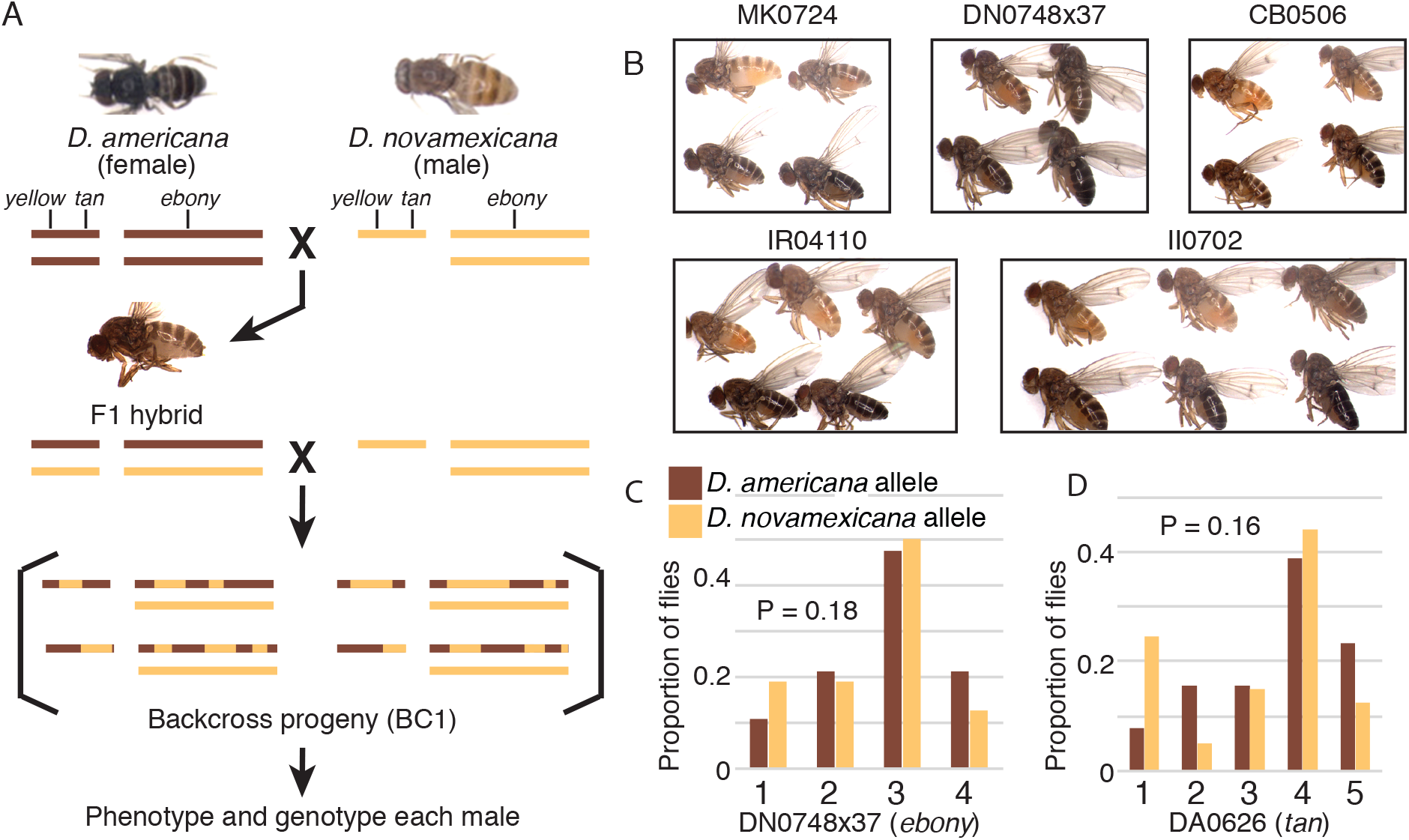
Genetic analysis of pigmentation differences between *D. novamexicana* and strains of *D. americana*. (A) Schematics show chromosomal content of *D. americana* and *D. novamexicana* parental strains, F_1_ hybrids, and examples of potential backcross progeny produced by crossing an F_1_ hybrid female back to *D. novamexicana*, with all autosomes represented as a single bar. Approximate locations of the *yellow* and *tan* genes on the X chromosome (Muller Element A) as well as the *ebony* gene on chromosome 2 (Muller element E) are also shown. Dorsal images of *D. novamexicana* (strain N14) and *D. americana* (strain CB0522) as well as the lateral image of a F_1_ hybrid shown were taken at different times from each other and images shown in panel B. Color adjustments have been made to reproduce relative pigmentation of these three genotypes, but these images should not be quantitatively compared to each other or images in panel B. (B) Representative flies from each of the 4 to 6 pigmentation classes identified for five strains of *D. americana* are shown, arranged from lightest (top left) to darkest (bottom right) in each box. A lateral view is shown for all flies and images within a box were collected under comparable conditions. (C, D) The proportion of male backcross flies in each pigmentation class carrying a *D. americana* (brown) or *D. novamexicana* (yellow) allele of *ebony* (C) or *tan* (D) inherited from their F_1_ hybrid mother is shown for backcrosses with two strains of *D. americana*: DN0748×37 (C) and DA0626 (D). These two examples are the only cases where no statistically significant difference in body color was detected for flies inheriting the *D. americana* or *D. novamexicana* alleles of *ebony* or *tan*. Phenotypic distributions are shown for *yellow*, *ebony*, and *tan* genotypes for all strains of *D. americana* in Supplementary Figures 3, 4 and 5, respectively. Note that borderline evidence of functional similarity for *tan* alleles was also observed between *D. novamexicana* and five other strains of *D. americana* (Supplementary Figure 5). None of the *D. americana* strains showed evidence of functional differences from *D. novamexicana* for alleles of the *yellow* gene (Supplementary Figure 3). Genotyping data for all three genes is provided as Supplementary Table 4, and results of the statistical tests are provided as Supplementary Table 5.

Consistent with prior descriptions of backcross populations between *D. americana* and *D. novamexicana* (Wittkopp *et al.* 2003; 2009), body color did not vary continuously within the BC_1_ populations. Rather, a limited number of distinct pigmentation categories were observed in each cross. The number of pigmentation classes ranged from four to eight among backcross populations produced by different strains; examples of pigmentation classes for five strains are shown in Figure 2B. The lightest (most yellow) body color phenotype in each backcross was assigned to category 1, with subsequent category numbers corresponding to progressively darker pigmentation.

To test for functional divergence of *ebony*, *tan*, or *yellow* alleles between *D. novamexicana* and each strain of *D. americana*, we calculated the difference in mean pigmentation score between flies that inherited the *D. americana* or *D. novamexicana* allele of each gene from their mother. For each gene and each BC_1_ population, the statistical significance of the pigmentation difference was determined by comparing it to a distribution of differences observed in 10,000 permuted datasets in which the genotypes were shuffled relative to the phenotypes. A false discovery rate correction for multiple tests (Benjamini & Hochberg 1995) was then applied, and an adjusted p-value cut-off of 0.05 was used to assess statistical significance. That is, tests with P < 0.05 were interpreted as evidence of functionally divergent alleles between *D. novamexicana* and the *D. americana* strain tested, whereas tests with P ≥ 0.05 were taken as evidence that the *D. novamexicana* and *D. americana* alleles were functionally equivalent. As expected, *yellow* alleles of *D. americana* and *D. novamexicana* appeared to be functionally equivalent for all strains tested (P > 0.14 in all cases; Supplementary Table 5; Supplementary Figure 3), further supporting the observation that *yellow* does not contribute to pigmentation divergence between these two species.

For *ebony*, all but one strain of *D. americana* tested showed evidence of functional divergence between *D. americana* and *D. novamexicana* (Supplementary Table 5; Supplementary Figure 4). This one exception (strain DN0748×37, Figure 2C) had a p-value of 0.18, suggesting that the *ebony* allele in this strain is functionally equivalent to the *D. novamexicana ebony* allele. Like the DN2 strain originally found to carry a *D. novamexicana-*like *ebony* allele, the DN0748×37 strain was collected from Duncan, Nebraska, but it was collected seven years later than the DN2 strain and did not share as much sequence similarity with the *D. novamexicana* allele as the DN2 allele (Figure 1D, Supplementary Figure 1). These observations suggest that more than one allele of *ebony* similar to *D. novamexicana* in function is segregating in the Duncan, Nebraska population. This population is located near the western edge of *D. americana*’s range (Figure 1C) and has some of the lightest pigmentation observed in *D. americana* (Wittkopp *et al.* 2011).

For *tan*, one strain of *D. americana* (DA0626) showed evidence of being functionally equivalent to the *D. novamexicana* allele (P = 0.16, Figure 2D, Supplementary Table 4). This strain was not any more similar in sequence to the *D. novamexicana tan* allele than other alleles of *D. americana* that showed evidence of functional divergence (Figure 1E, Supplementary Figure 2). Five other *D. americana* strains showed marginal evidence of being functionally equivalent to the *D. novamexicana* allele (P-values = 0.05 or 0.06, Supplementary Figure 5, Supplementary Table 5). With all other strains showing P-values < 0.0001 (Supplementary Table 5), these five alleles are interpreted as being at least functionally distinct from the majority of *D. americana tan* alleles, if not equivalent to the *D. novamexicana tan* allele. Two of these five alleles were found in strains collected from the same population (SC0708, SC0718) near the western edge of the species range; however, the other three alleles (II0710, G9647, FP9918, DA0626) as well as the DA0626 allele were found in strains isolated from populations spread throughout the species range (Figure 1C).

The frequency and geographic distribution of *ebony* and *tan* alleles similar in function to their *D. novamexicana* orthologs again suggests that they are unlikely to be primarily responsible for the pigmentation cline.

### Testing for associations between pigmentation and variation in *ebony* and *tan*

Although we found few alleles with sequence and/or function equivalent to *D. novamexicana* segregating within *D. americana*, other alleles of *tan* and/or *ebony* might still contribute to pigmentation diversity within *D. americana*. To explore this possibility, we tested whether any of the segregating sites sampled in *tan* (Supplementary Table 6) or *ebony* (Supplementary Table 7) for our phylogenetic analysis showed a significant association with estimates of pigmentation for each strain (Supplementary Table 8). Specifically, we used a general linear model to test each variable site with a minor allele present in at least five strains (excluding sites with indels) for a statistically significant association with pigmentation. For *ebony*, the region sampled started in exon 5 and extended into exon 8, with no statistically significant associations observed (Figure 3A). Because prior work suggests that the functional difference between *D. americana* and *D. novamexicana ebony* alleles affects *cis*-regulation (Cooley *et al.* 2012), it is perhaps not surprising that this region, consisting mainly of coding sequences, does not harbor associated variants. We thought it possible, however, that we might have seen an association with these sites due to linkage disequilibrium with a variant outside this region because *ebony* is located in a region of the genome inverted between *D. novamexicana* and most strains of *D. americana* (Wittkopp:2003bn; Wittkopp *et al.* 2009). For *tan*, prior work has mapped functionally divergent sites to intron 1 (Wittkopp *et al.* 2009), suggesting that the region sampled is much more likely to harbor variants that might correlate with pigmentation. Nonetheless, we also observed no statistically significant associations between body color and variants in this region segregating within *D. americana* (Figure 3B).

**Figure 3.**
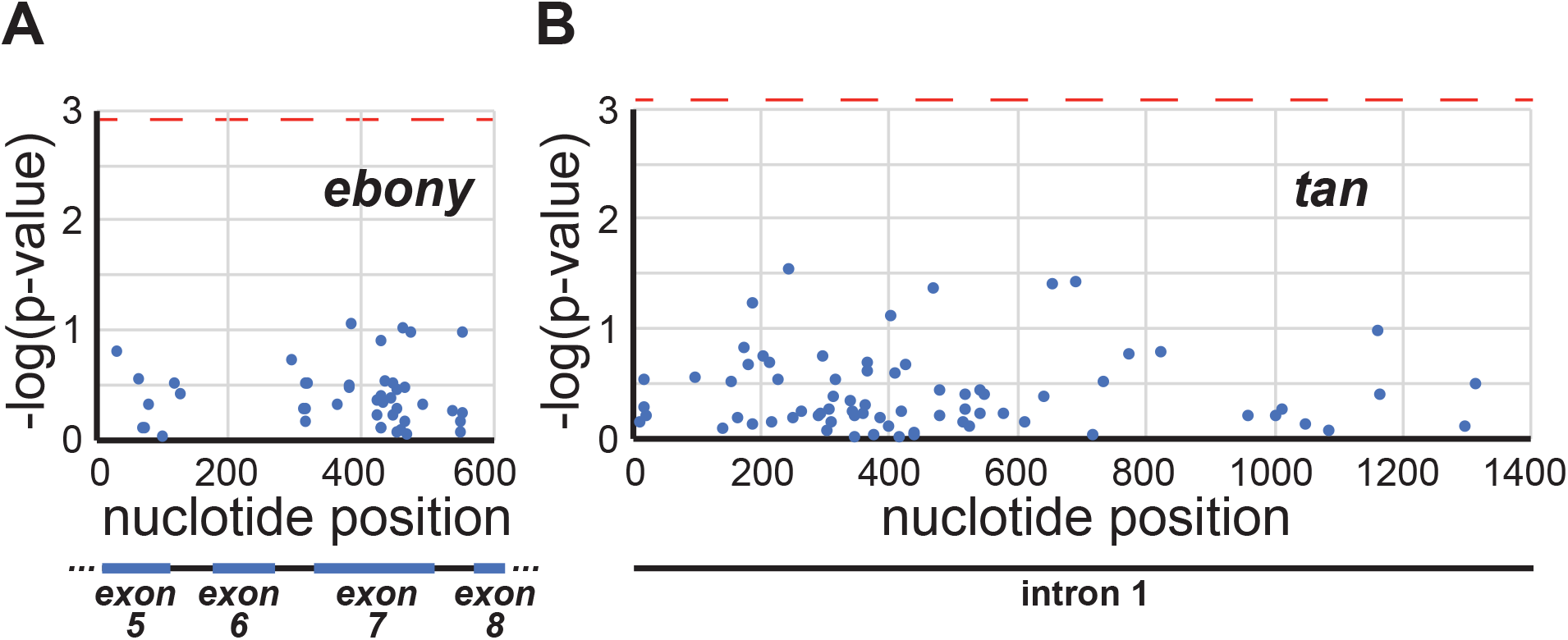
Variable sites sampled in *tan* and *ebony* are not significantly associated with pigmentation in *D. americana*. Statistical significance of an association between body color and the nucleotide present at variable sites in the *D. americana ebony* (A) and *tan* (B) regions sequenced are shown, reported as −log(p-value) from the general linear model described in Methods. Red dotted lines show threshold used to assess statistical significance. Schematics shown below each plot indicate the location of intronic and exons regions in the *ebony* (A) and *tan* (B) sequences analyzed. Body color data used provided as Supplementary Table 2. Genotype data used provided as Supplementary Table 6 for *tan* and Supplementary Table 7 for *ebony*. Results of the general linear models are provided as Supplementary Table 8.

### Genetic heterogeneity underlying body color variation in D. americana

With none of our analyses linking variation in *ebony* and/or *tan* to clinal variation in *D. americana* body color, we sought to further investigate its genetic architecture by examining the phenotypic distributions of males in the 51 backcross populations. Because all 51 strains were crossed and then backcrossed to the same strain of *D. novamexicana*, differences in the distribution of pigmentation phenotypes observed among these BC_1_ populations must be due to genetic differences among the strains of *D. americana*. For example, differences in the number of phenotypic classes observed among the BC_1_ populations indicate that different strains of *D. americana* harbor different numbers of loci with effects on pigmentation distinct from the *D. novamexicana* alleles. Assuming basic Mendelian segregation, one locus with a divergent allele affecting pigmentation is expected to cause two distinct pigmentation phenotypes in the backcross population, whereas two loci with divergent alleles are expected to cause up to four distinct pigmentation phenotypes, and three loci with divergent alleles could cause up to eight distinct phenotypes. Differences in the BC_1_ pigmentation phenotypes and/or number of pigmentation categories are also expected to result from variation among the *D. americana* strains in the identity of loci and/or allelic variation at loci.

To compare the distributions of BC_1_ phenotypes among strains, we first converted the strain-specific pigmentation categories to a standardized set of pigmentation categories. We did this by comparing representative images of flies from each strain-specific category to each other and sorting the images with the most similar pigmentation into the same category. This process resulted in 8 categories. After translating the numbers of flies from the strain-specific categories to the standardized categories (Supplementary Table 3), we examined the distribution of flies among pigmentation classes for all of the strains. We found that the number of pigmentation categories in the BC_1_ population ranged from 4 (e.g., BU0624) to 8 (WS0712) among the strains (Supplementary Table 3; Figure 4A), indicating that the number of loci harboring variation affecting pigmentation is variable within *D. americana*. In addition, even for strains that produced the same number of phenotypic classes in the backcross population, differences were observed in the specific pigmentation phenotypes of each class, indicating that there are also differences in the specific loci or alleles affecting pigmentation between strains. An example of this can be seen by comparing strains BU0624 and PM9936: both strains produced backcross populations with 4 pigmentation classes, but flies with light pigmentation were common in the BU0624 backcross and nonexistent in the PM9936 backcross (Figure 4A).

**Figure 4.**
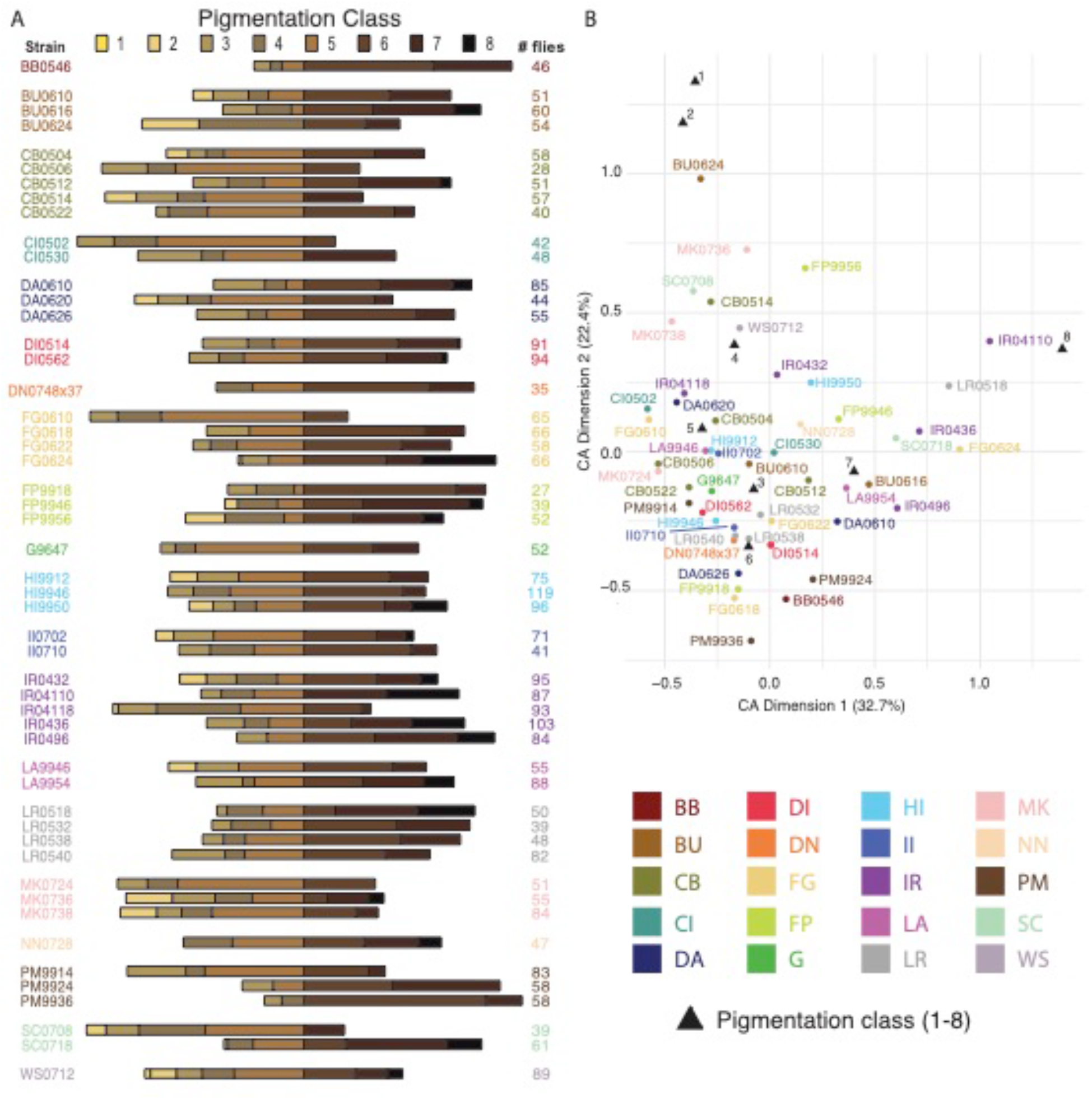
Distributions of backcross phenotypes indicate diversity in number and effects of loci affecting pigmentation. (A) The relative proportion of male backcross progeny in each of eight standardized pigmentation classes (Supplementary Table 3) is shown for each *D. americana* strain. Pigmentation classes are indicated by the color of the bar ranging from the lightest (yellow, class 1) to the darkest (black, class 8), with a longer bar indicating a greater proportion of the backcross population. Bars are aligned vertically at the transition between pigmentation classes 5 and 6. Strains are clustered by collection site, with each strain derived from the same collection site shown in the same color. The total number of male backcross progeny scored for each strain is shown to the right of each distribution. Note the differences in distributions not only between, but also within, collection sites. For example, strains producing very different distributions of backcross progeny were isolated from the FG, IR, and SC collection sites. (B) Results from a correspondence analysis (CA) used to compare the distribution of backcross pigmentation phenotypes among strains are shown, plotted with colored circles according to their values on the first two axes of variation: CA dimension 1, which explained 32.7% of the variation and CA dimension 2, which explained 22.4% of the variation. Strains shown with the same color were derived from the same collection site. The relative placement of pigmentation classes 1-8 on these two axes is also shown with black triangles for comparison. Note that, for example, strain IR4110, which had most backcross progeny with the darkest body color is located close to the triangle representing the darkest pigmentation class (class 8). Similarly, BU0624, the strain that produced the most lightly pigmented backcross progeny, is located close to the triangles representing the lightest pigmentation classes (class 1 and 2). The lack of visual clustering for strains derived from the same collection site is consistent with our statistical test showing strains from the same collection site were no more likely to be located close to each other in this CA space than strains from different collection sites.

Finally, we asked whether loci affecting pigmentation were more likely to be more similar for strains isolated from the same population than from different populations. Despite evidence of extensive gene flow within *D. americana* (Schäfer *et al.* 2006; Morales-Hojas *et al.* 2008; Fonseca *et al.* 2013), we expected this might be true for loci affecting pigmentation because of the longitudinal cline previously observed for body color (Wittkopp *et al.* 2011). That is, if natural selection is favoring different pigmentation phenotypes in different populations, we might expect to see more genetic similarity for loci affecting pigmentation within than between populations. Inspecting the number of backcross pigmentation categories for strains derived from the same collection site, however, already suggests this might not be so: the three strains isolated from the MK population produced backcross progeny with 4, 6, and 7 distinct pigmentation phenotypes.

To further compare the backcross phenotypes, we used correspondence analysis (CA) to reduce the dimensionality of the BC_1_ phenotypic distributions. This method is similar to principal components analysis (PCA), but for categorical data. The first two dimensions of the correspondence analysis (comparable to the first two principle components in a PCA) captured 55.1% of the variation among strains. As seen by the overlaid pigmentation categories in Figure 4B, dimension 1 discriminates most strongly between strains that do and do not produce many backcross progeny with the darkest pigmentation (categories 7 and 8). Dimension 2, by contrast, discriminates most strongly between strains that do and do not produce many backcross progeny with the lightest pigmentation (categories 1 and 2) (Figure 4B). The lack of visible clustering for strains isolated from the same collection site again suggests that flies in the same population might not be more likely to have similar loci affecting pigmentation than flies from different populations. Indeed, Euclidean distances in this CA dimension 1 and 2 space were similar for the 110 pairs of strains from the same collection site and the 2440 pairs of strains that were from different collection sites (mean distance for pair from same collection site = 0.68; mean distance for pairs from different collection sites = 0.65; t-test, p-value = 0.45).

## Discussion

In this study, we tested the hypothesis that *D. novamexicana*-like alleles of *ebony* and/or *tan* are driving the longitudinal pigmentation cline seen in *D. americana* (Wittkopp *et al.* 2009; 2011; Cooley *et al.* 2012). We found no support for this hypothesis: *D. novamexicana*-like alleles of these genes segregating in *D. americana* - identified based on either sequence or function - were too rare to account for the cline. Other alleles of *tan* and/or *ebony* might contribute to pigmentation variation within *D. americana*, but we found no statistically significant association between body color and any of the variable sites in *tan* or *ebony* tested. Rather, genetic analysis indicated that differences in the number of loci and/or allelic effects of loci affecting pigmentation are common both within and among populations, suggesting genetic heterogeneity despite locally adapted pigmentation. Below, we discuss the implications of these findings, focusing on possible sources of pigmentation variation in *D. americana*, the complexity of its genetic architecture, and how this pigmentation cline might persist in the face of ongoing gene flow.

In other Drosophila species, differences in body pigmentation segregating within a species have been shown to be associated with variable sites in pigmentation genes, including *ebony* (Pool & Aquadro 2007; Takahashi *et al.* 2007; Rebeiz *et al.* 2009; Telonis-Scott *et al.* 2011; Takahashi & Takano-Shimizu 2011; Bastide *et al.* 2013; Johnson *et al.* 2015; Miyagi *et al.* 2015; Telonis-Scott & Hoffmann 2018) and *tan* (Bastide *et al.* 2013; Yassin *et al.* 2016; Endler *et al.* 2016). Despite the lack of associations observed in the current study, we still think it likely that variation in *ebony*, *tan*, and/or other pigmentation genes also contribute to pigmentation variation within *D. americana*. We tested for associations between pigmentation and variable sites in *ebony* and *tan* using ~100 strains each, but larger sample sizes would provide greater power to detect variants with small effects. In addition, we only tested segregating sites in the first intron of *tan* and in a region starting in exon 5 and ending in exon 8 for *ebony*. Because linkage disequilibrium in *D. americana* decays quickly within these genes (often disappearing within ~50 bp) (Wittkopp *et al.* 2009; 2011), it is unlikely that the sites tested would detect functional variants outside of these regions; variable sites in other regions of *tan* and/or *ebony* might be found to be associated with *D. americana* body color in future studies.

Association studies can also fail to identify genes contributing to trait variation when there is genetic heterogeneity (i.e., multiple genotypes giving rise to the same phenotype) (Korte & Farlow 2013; Manchia *et al.* 2013). Genetic heterogeneity is expected to be more common for polygenic than single-gene traits, but even when there is only one gene controlling a trait, allelic heterogeneity (multiple alleles with the same phenotypic effects) can still obscure associations with the gene (Savolainen *et al.* 2013). Our genetic analysis provides two lines of evidence for such heterogeneity underlying pigmentation variation in *D. americana*. First, for *tan*, we identified six *D. americana* alleles showing at least marginal evidence of similarity between *D. americana* and *D. novamexicana,* indicating that they lighten pigmentation more than other *D. americana tan* alleles, but these alleles were derived from five different collection sites in four different states (Alabama, Arkansas, Indiana, and Missouri) and in only one case were two of these alleles sampled from the same collection site. This finding suggests that the similar pigmentation of strains collected from these sites exists despite differences in the pigmentation alleles they carry. A similar pattern was reported previously for *D. americana* when a *D. novamexicana*-like *ebony* allele causing lighter pigmentation was found to be present in one of three strains with similar pigmentation derived from Duncan, Nebraska (Wittkopp *et al.* 2009). Indeed, these *D. novamexicana*-like *tan* and *ebony* alleles found segregating in *D. americana* provide an excellent example of how genetic heterogeneity can work: because *ebony* and *tan* encode enzymes catalyzing opposite directions of a reversible biochemical reaction (True *et al.* 2005), alleles increasing activity of *ebony* and decreasing activity of *tan* (or vice versa) can have equivalent effects on pigmentation (Figure 1B, (Wittkopp *et al.* 2009).

Our phenotypic analysis of backcross populations from 51 strains of *D. americana* from 20 collection sites provides the second line of evidence for genetic heterogeneity underlying clinally varying pigmentation in *D. americana*. In the absence of genetic heterogeneity, two strains derived from the same population with the same phenotype are expected to carry the same pigmentation alleles. If true, crossing and backcrossing these strains of *D. americana* to *D. novamexicana* should produce the same distributions of pigmentation phenotypes. We found, however, that backcross populations often showed differences in the number of distinct pigmentation classes, the body color of each pigmentation class, and/or the relative abundance of flies with different body colors, even when strains were derived from the same collection site. These data are consistent with genetic heterogeneity in which multiple combinations of genes and/or alleles underlie similar pigmentation phenotypes within a population as well as diversity in pigmentation among locations. Similar genetic heterogeneity has previously been described for mate choice in *Drosophila pseudoobscura* (Barnwell & Noor 2008), gene expression in yeast (Metzger & Wittkopp 2019), timing of bud set in Scots pine trees (Kujala *et al.* 2017), flowering time in maize (Buckler *et al.* 2009), and human diseases (McClellan & King 2010). It has also been reported more broadly for convergent phenotypes that evolved in more genetically isolated populations, including adaptation of humans to high-altitudes (Jeong & Di Rienzo 2014), lighter skin color in East Asian and European peoples (Norton *et al.* 2007), and adaptation to highlands in maize (Takuno *et al.* 2015). Nonetheless, we think that the extent of genetic heterogeneity underlying variation in quantitative traits is generally underestimated - especially within a population or among populations connected by extensive gene flow - because of the reliance on association mapping for finding loci responsible for trait variation and the rarity of studies using biparental quantitative trait locus (QTL) mapping to analyze multiple genotypes from the same population with similar phenotypes.

How might this genetic complexity be maintained despite selection favoring a particular phenotype at a particular location? The extensive gene flow seen throughout *D. americana* (Schäfer *et al.* 2006; Morales-Hojas *et al.* 2008; Wittkopp *et al.* 2011; Fonseca *et al.* 2013) is likely part of the answer. This gene flow moves alleles among populations, making it difficult for a population to fix the most adaptive allele for each local environment (Savolainen *et al.* 2013). But there must also be sufficient genetic variation affecting pigmentation maintained in the species for this gene flow to cause genetic heterogeneity (Pritchard *et al.* 2010; Savolainen *et al.* 2013). *D. americana* harbors high levels of genetic variation generally (Fonseca *et al.* 2013), and selection for different pigmentation phenotypes in different locations should maintain diverse pigmentation alleles at the species level (Savolainen *et al.* 2013; Lee *et al.* 2016; Troth *et al.* 2018). The structure of the biochemical pathway controlling production of alternative pigments from a single, branched biochemical pathway (Massey & Wittkopp 2016) might also contribute to standing genetic variation because it allows changes in the activity of multiple genes to have similar effects on pigmentation (Wittkopp *et al.* 2009). Ultimately, however, selection acting on this standing genetic variation must be favoring different pigmentation phenotypes in different locations to maintain the cline (Kawecki & Ebert 2004; Savolainen *et al.* 2013). Assortative mating, in which individuals with similar body color are more likely to mate with each other than individuals with different body color, could also contribute to the *D. americana* pigmentation cline. Although evidence of assortative mating for body color is rare in Drosophila species, it has been observed in an Indian population of *D. melanogaster*, with darker individuals more likely to mate with each other in cold, dry weather and lighter individuals more likely to mate with each other when it is hot or humid (Dev *et al.* 2013). Disentangling the relative contributions of these different evolutionary and molecular processes to the formation and maintenance of the *D. americana* body color cline will require much more extensive, interdisciplinary studies.

## Acknowledgements

We thank Bryant McAllister for sharing strains of *D. americana* that his lab established from field collections, the Drosophila Species Stock Center for maintaining and supplying strains of *D. novamexicana* and *D. americana*, and members of the Wittkopp lab (especially Henry Ertl, Mark Hill, Petra Vande Zande, Abigail Lamb and Molly Hirst) for helpful discussions and feed-back on this work. Funding for this project was provided by the National Institutes of Health (F32GM087928 to AC; R35GM118073 and R01GM08973 to PJW) and the National Science Foundation (DEB-0640485 to PJW).

## Data Accessibility

Sequences described in Supplementary Files 1 and 2 are also available in NCBI PopSet with GenBank accession numbers: MT350927 - MT351036 for *ebony* and MT350824 - MT350926 for *tan*. All other data and code are included in the manuscript as supplementary tables and files.

## Author Contributions

PJW and LLS designed the research. LLS and WNM quantified pigmentation and performed the genetic analysis. AMC, DCY, AJ, and PJW collected and analyzed sequence data. PJW performed the statistical analysis and constructed figures, with assistance from LLS and WNM. PJW wrote the paper, with input from LLS, WNM, and AJ and final editing by all authors.

## Supplemental Information

**Supplementary Figure 1.**
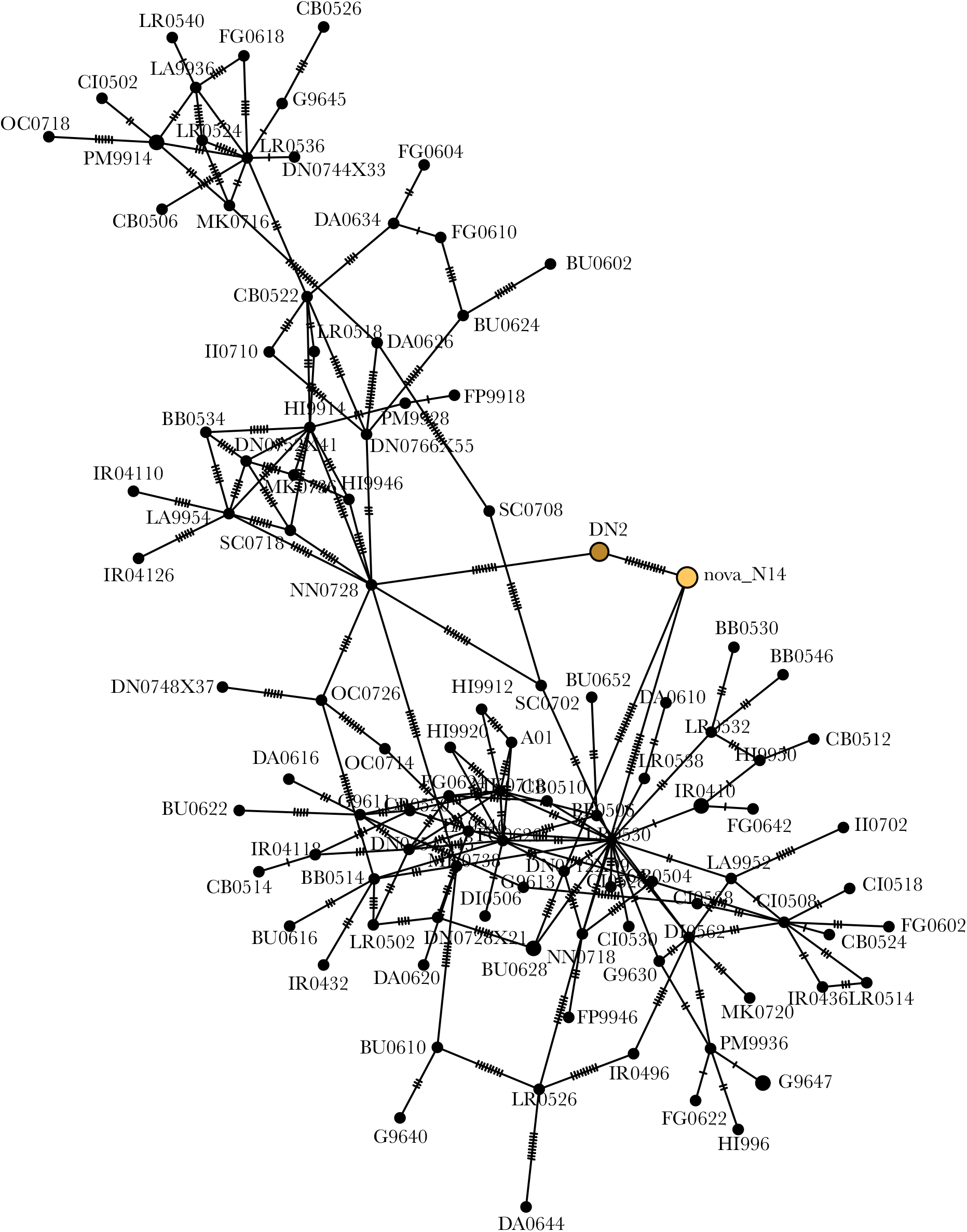
Haplotype network for *ebony*. A Median Spanning Network built from the same *ebony* sequences used to construct the phylogenetic tree shown in Figure 1D is shown. Note that the DN2 allele from *D. americana* previously shown to share similarity in sequence and function with *D. novamexicana* (brown) is most similar to the *D. novamexicana* (“nova_N14”) allele (yellow).

**Supplementary Figure 2.**
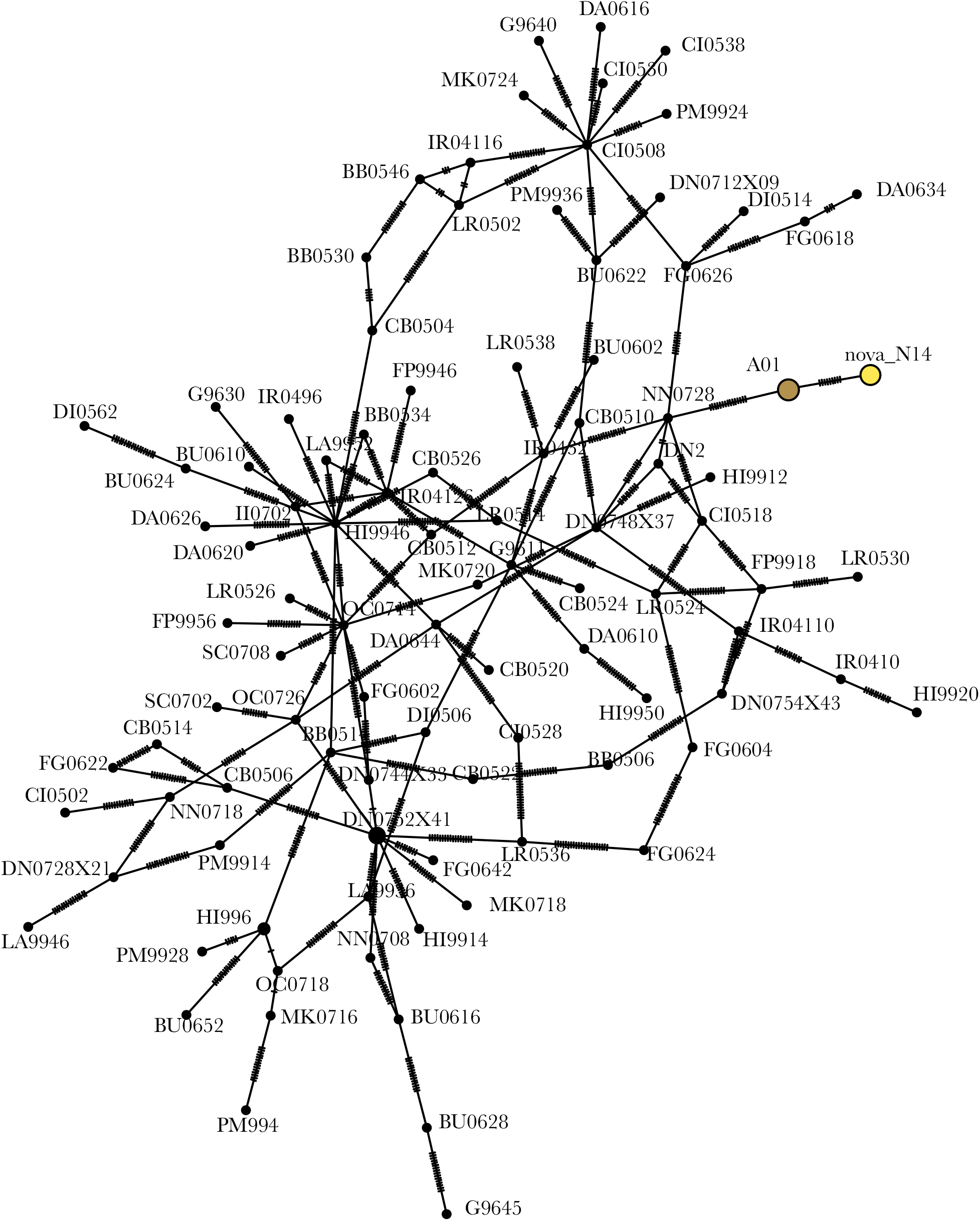
Haplotype network for *tan*. A Median Spanning Network built from the same *tan* sequences used to construct the phylogenetic tree shown in Figure 1E is shown. Note that the A01 allele from *D. americana* previously shown to share similarity in sequence and function with *D. novamexicana* (brown) is most similar to the *D. novamexicana* (“nova_N14”) allele (yellow).

**Supplementary Figure 3.**
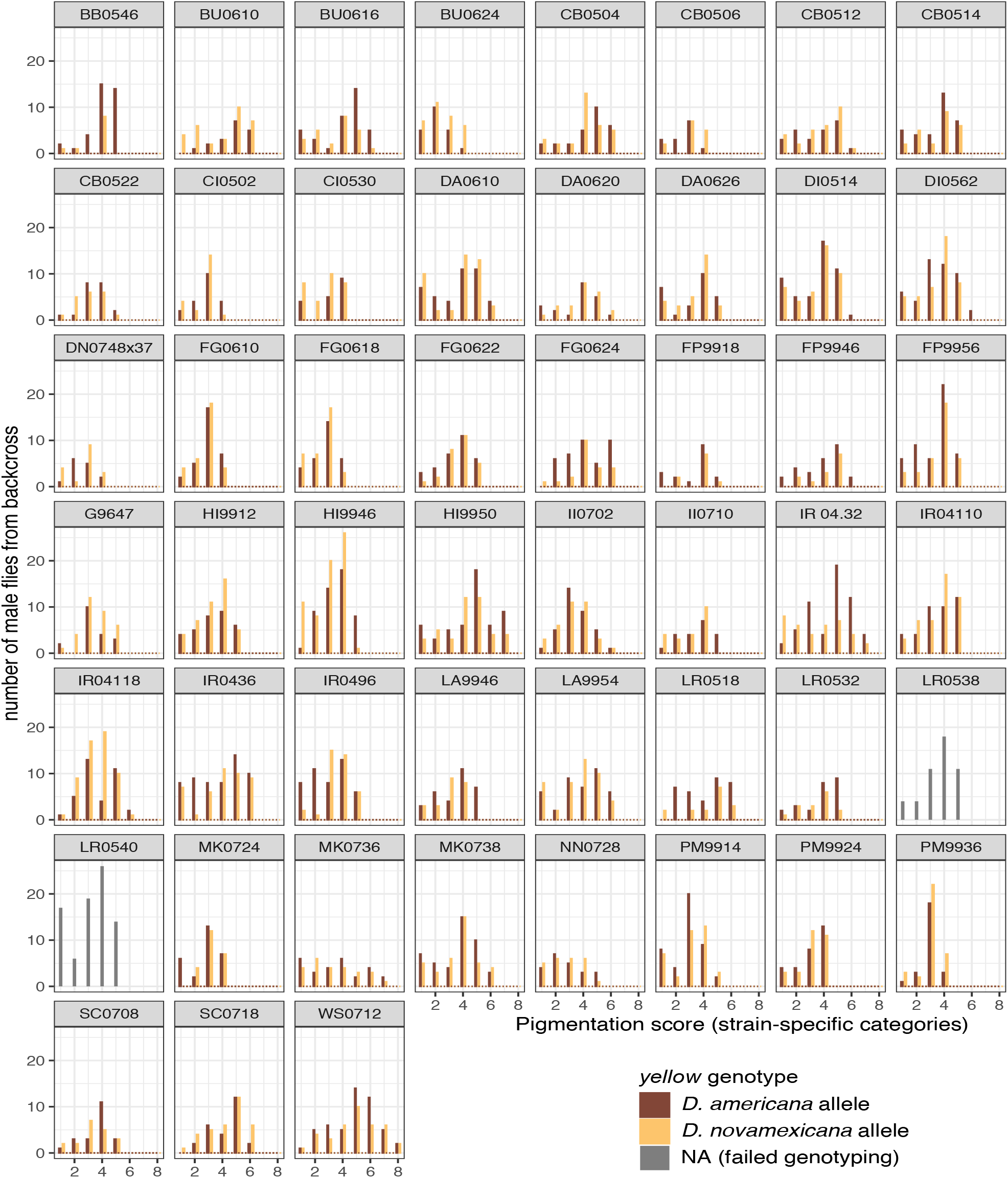
Testing for function divergence of *yellow*. Distributions of pigmentation phenotypes for backcross progeny inheriting the *D. americana* (brown) or *D. novamexicana* allele (yellow) of the *yellow* gene from their F_1_ hybrid mother are shown for each strain of *D. americana* tested, with the stain name shown at the top of each panel. Numbers of males in each pigmentation class are shown rather than proportions to communicate sample sizes. Grey bars are shown for strains LR0538 and LR0540 because the *yellow* genotyping assay failed for all flies, presumably because of sequence differences in these *yellow* alleles.

**Supplementary Figure 4.**
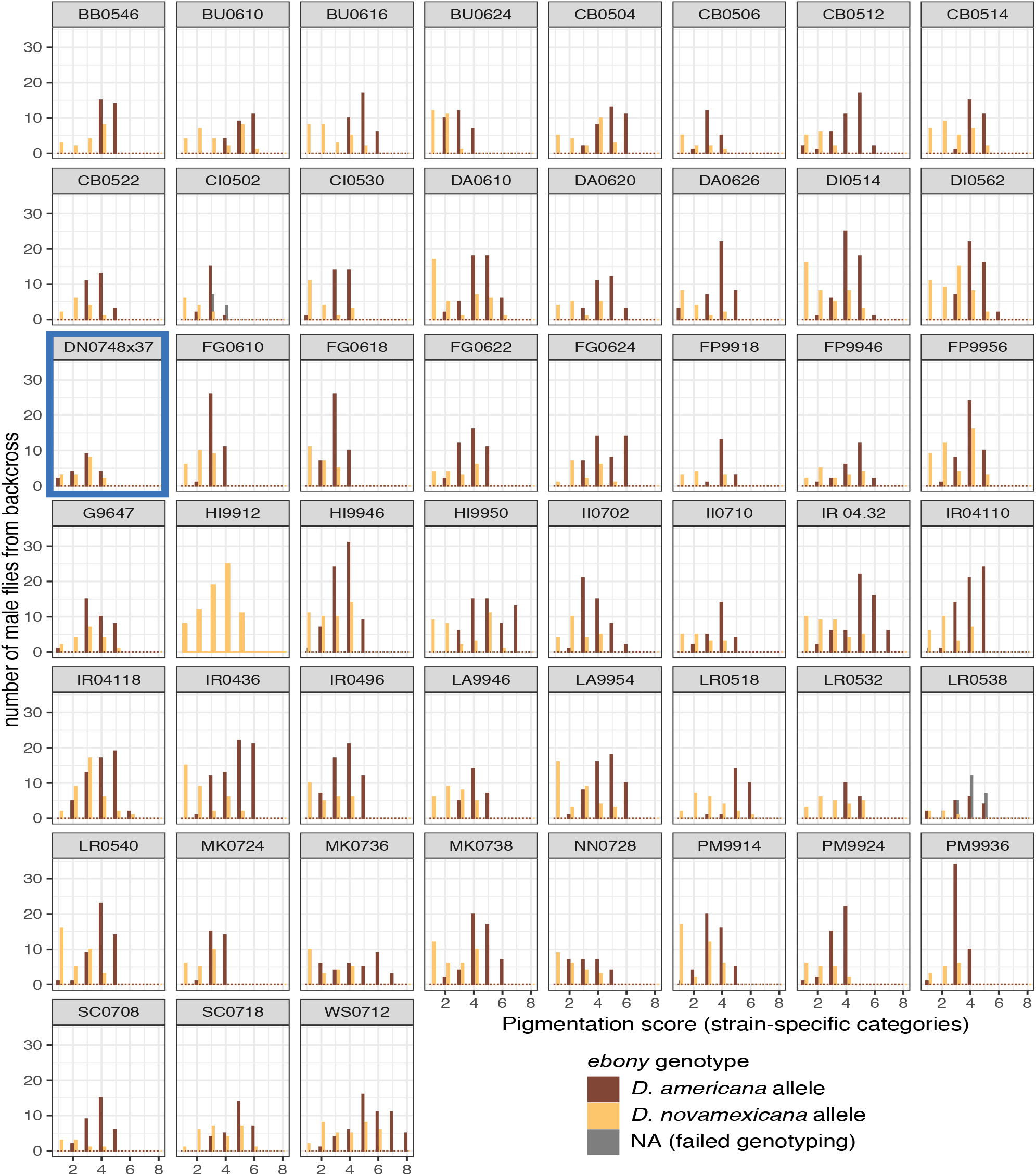
Testing for function divergence of *ebony*. Distributions of pigmentation phenotypes for backcross progeny inheriting the *D. americana* (brown) or *D. novamexicana* allele (yellow) of the *ebony* gene from their F_1_ hybrid mother are shown for each strain of *D. americana* tested, with the stain name shown at the top of each panel. Numbers of males in each pigmentation class are shown rather than proportions to communicate sample sizes. Grey bars indicate samples with failed genotyping reactions, which were most common for *ebony* with flies from strain LR0538. The genotyping assay seemed to fail to differentiate alleles in the backcross with strain HI9912. Dark blue box indicates no significant difference between *D. americana* and *D. novamexicana* alleles.

**Supplementary Figure 5.**
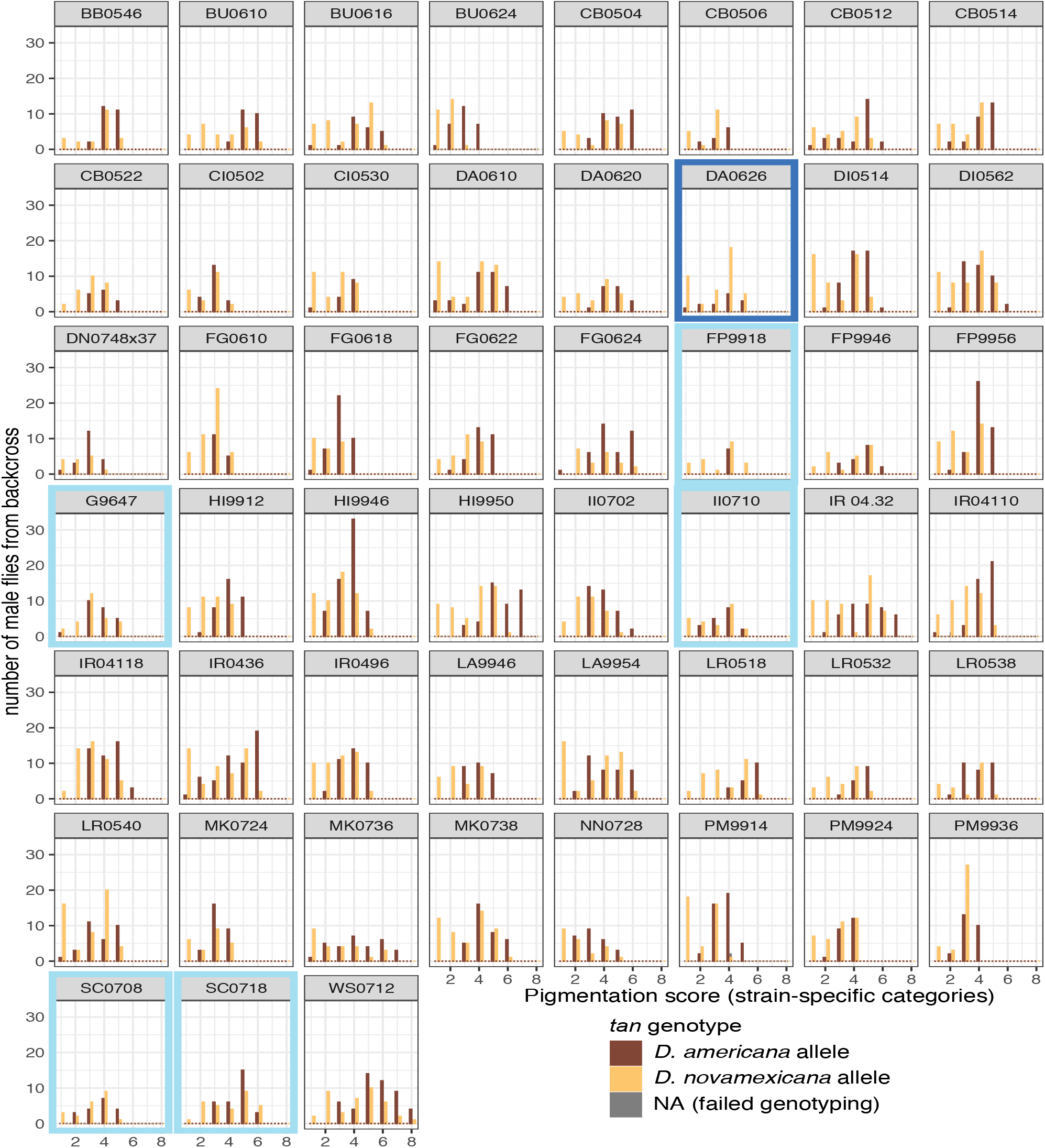
Testing for function divergence of *tan*. Distributions of pigmentation phenotypes for backcross progeny inheriting the *D. americana* (brown) or *D. novamexicana* allele (yellow) of the *tan* gene from their F_1_ hybrid mother are shown for each strain of *D. americana* tested, with the stain name shown at the top of each panel. Numbers of males in each pigmentation class are shown rather than proportions to communicate sample sizes. Grey bars indicate samples with failed genotyping reactions, of which very few were observed for *ebony*. Dark blue box indicates no significant difference between *D. americana* and *D. novamexicana* alleles. Light blue boxes indicate marginal evidence of equivalent alleles (P-values = 0.05 or 0.06).

### Supplementary Tables

**Supplementary Table 1.** Summary of strains used for sequence analysis and/or functional testing, including details of sites where their progenitors were collected.

**Supplementary Table 2.** Median pigmentation measure for each fly sampled from each strain of *D. americana* obtained using a custom-built fiber optic probe to measure light reflected off the fly’s abdominal cuticle.

**Supplementary Table 3.** Standardization of pigmentation classes among all backcrosses.

**Supplementary Table 4.** *yellow*, *tan*, and *ebony* genotypes for male progeny of F_1_ hybrids backcrossed to *D. novamexicana*.

**Supplementary Table 5.** Results of permutation tests used to identify functional differences between *D. americana* and *D. novamexicana* alleles of *yellow*, *tan*, and *ebony*.

**Supplementary Table 6.** Genotypes of sites in *tan* used to test for an association with body pigmentation.

**Supplementary Table 7.** Genotypes of sites in *ebony* used to test for an association with body pigmentation.

**Supplementary Table 8.** Results from general linear models used to test for associations between body pigmentation and variable sites in *tan* and *ebony.*

### Supplementary Files

**Supplementary File 1.** FASTA format summary of *ebony* allele sequences analyzed.

**Supplementary File 2.** FASTA format summary of *tan* allele sequences analyzed.

**Supplementary File 3.** Text file containing R code used for all analyses presented in the manuscript.

## Notes

### Competing Interest Statement

The authors have declared no competing interest.

