## Supplementary File 3 for "Genetic architecture of a body color cline in *Drosophila americana*"

###################################
## The code below was used for analyses described in Sramkoski et al. 2020 titled
## "Genetic architecture of a body color cline within Drosophila americana"
###########################

#######################
## This file contains code for the following parts of the work:
## [1] Estimating pigmentation for each strain
## [2] Plotting distributions of BC1 pigmentation phenotypes (standardized) by strain and genotype
## [3] Testing for relationship between tan, ebony, and yellow genotypes with pigmentation in BC1
## [4] Testing for association between variable sites in tan and ebony with pigmentation
## [5] Correspondence analysis
## code by Patricia J. Wittkopp
#######################

##########################
## [1] The code below was used to estimate pigmentation for strains from the following populations:
## BB, BU, CB, CI, DA, DI, FG, FP, G, HI, IR, LA, LR, PM.
## LSMEANS estimating pigmentation for strains from DN, II, MK, NN, OC, SC, and WS populations
## were taken from Wittkopp et al. (2011) Heredity
## More positive values = lighter pigmentation
## Summary Table of median pigmentation per fly provided as Supplementary Table 2

library(dplyr)
library(lme4)
library(lsmeans)

library(foreach)

#import data
library(readxl)
americana_lines_pigmentation_round_2_Lisa <- read_excel("americana lines pigmentation round 2 Lisa.xlsx")

#calculate median for each fly scored
med_pig<-pig %>%
group_by(Strain, fly_number) %>%
summarize(Median=median(pigment_score))

#count number of flies scored per strain
N_pig<-med_pig %>%
group_by(Strain) %>%
summarize(N=n())

##For lines used, minimum number of flies scored was 6 (LR0532)
min(N_pig$N)
max(N_pig$N)


#Export table of medians by line
write.table(med_pig, file="pigmentation_medians.txt", sep="\t")

#Note: Population designations added to each entry outside of R
#Reimport medians (provided in Supplementary Table 2)
pigmentation_medians <- read.delim("~/pigmentation_medians.txt")

#fit GLM and extract lsmeans

exp.model<-lmer(Median~Strain+(1|fly_number), data=pigmentation_medians)
lsmeans<-lsmeans(exp.model,pairwise~Strain)

## median of 5 replicate pigmentation measures for each fly.
## LSMeans from fitting medians from 6-20 replicate flies to the following model:
## median = Strain(fixed) + fly (random).  Some lines (see above) using lsmeans from HDY paper.
## more positive score = lighter pigmentation


##############################
## [2] Plotting distributions of backcross phenotypes by strain using standarized pigmentation scale
## used to make Supplementary figures 3, 4, and 5

amer_genos$ebony<-as.factor(amer_genos$ebony)
ggplot(amer_genos,aes(x=Phenotype,group=ebony,fill=ebony))+
geom_histogram(position="dodge",binwidth=0.25)+theme_bw()+
facet_wrap( ~ `Line name`)

amer_genos$tan<-as.factor(amer_genos$tan)
ggplot(amer_genos,aes(x=Phenotype,group=tan,fill=tan))+
geom_histogram(position="dodge",binwidth=0.25)+theme_bw()+
facet_wrap( ~ `Line name`)

amer_genos$yellow<-as.factor(amer_genos$yellow)
ggplot(amer_genos,aes(x=Phenotype,group=yellow,fill=yellow))+
geom_histogram(position="dodge",binwidth=0.25)+theme_bw()+
facet_wrap( ~ `Line name`)


#############################
## [3] Code below tests for differences in the effect of D. americana and D. novamexicana alleles
## of ebony, tan, and yellow on body color (pigmentation). Prior work shows that ebony and tan
## have functionally diverged between D. novamexicana and most lines of D. americana, whereas
## yellow alleles have previously been found to be functionally equivalent (Wittkopp et al. 2009 Science).
## There are three separate blocks of code, one for each of the three genes tested.
## In all cases, input data used ("amer_genos.xlsx") is the full file provided as Supplementary Table 4.

################
#ebony

library(readxl)
amer_genos <- read_excel("amer_genos.xlsx")

amer_genos$yellow <- factor(amer_genos$yellow)
amer_genos$tan <- factor(amer_genos$tan)
amer_genos$ebony <- factor(amer_genos$ebony)
amer_genos$Line <- factor(amer_genos$Line)

#divide datafile by gene and remove flies with missing genotypes
E<-subset(amer_genos, select=c(Line, Phenotype, ebony))#pulls out data relevant for ebony
E_clean<-na.omit(E) #removes flies with missing genotype data for ebony
E_clean.split <- split(E_clean$Phenotype,E_clean$Line) #pulls out list of phenotypes for each line


#gets a list of Lines with ebony genotypes, n=51 for ebony
Lines<-unique(E_clean$Line)
Lines<-sort(Lines)
Lines<-as.character(Lines)


#summarize information about ebony genotypes/phenotypes for each line
LINE_SUMMARY<- function(x) {
    line<-x
    obs_diff<-mean(E_clean[["Phenotype"]] [which(E_clean[["Line"]]==x & E_clean[["ebony"]]=="0")]) - mean(E_clean[["Phenotype"]][which(E_clean[["Line"]]==x & E_clean[["ebony"]]=="1")])
    n_amer<-NROW(E_clean[["Phenotype"]] [which(E_clean[["Line"]]==x & E_clean[["ebony"]]=="0")])
    n_nova<-NROW(E_clean[["Phenotype"]][which(E_clean[["Line"]]==x & E_clean[["ebony"]]=="1")])
    n_tot<-NROW(E_clean[["Phenotype"]][which(E_clean[["Line"]]==x)])
    sum_amer<-sum(E_clean[["Phenotype"]] [which(E_clean[["Line"]]==x & E_clean[["ebony"]]=="0")])
    sum_nova<-sum(E_clean[["Phenotype"]][which(E_clean[["Line"]]==x & E_clean[["ebony"]]=="1")])
    sum_tot<-sum(E_clean[["Phenotype"]][which(E_clean[["Line"]]==x)])
    avg_amer<-sum_amer/n_amer
    avg_nova<-sum_nova/n_nova
    avg_tot<-sum_tot/n_tot
    obs_diff<-avg_amer - avg_nova
    
    results<- list(line, n_amer, n_nova, n_tot, sum_amer, sum_nova, sum_tot, avg_amer, avg_nova, avg_tot, obs_diff)
    return(results)
}

line_summary_data<-sapply(Lines, LINE_SUMMARY, simplify=TRUE, USE.NAMES=FALSE)
rownames(line_summary_data) <- c("Line", "n_amer", "n_nova", "n_tot", "sum_amer", "sum_nova", "sum_tot", "avg_amer", "avg_nova", "avg_tot", "obs_diff")
colnames(line_summary_data)<-Lines
write.table(line_summary_data, "./E_line_summary_data.txt", sep="\t")

line_obs_diff_E<-line_summary_data[11,] #summary of observed differences by line
counts<-line_summary_data[2,] #number of D. americana alleles by line


#Define permutation values and pvalue collection list
p_val<- rep(NA, length(Lines))
N.PERM=10000

#resample function
P_val_calc_E<-function(x,y,z,q) #x=E_clean.split (pigmentation scores by line), y = number of americana genos per line, z = lines, q=obs_diff
{
    perm_dist <- rep(NA,N.PERM) #resest list for permuted values of difference between amer and nova
    line<-z
    for (k in 1:N.PERM)
    {
        sum_perm_amer<-sum(sample(x,y, replace=FALSE, prob=NULL)) #sample without replacement, equal frequency
        mean_perm_amer<-sum_perm_amer/y
        obs_sum<-sum(x) #sum of pig score for line
        sample_size<-length(x) #sample size for line
        sum_perm_nova<-obs_sum-sum_perm_amer
        mean_perm_nova<-sum_perm_nova/(sample_size-y)
        perm_diff<-mean_perm_amer - mean_perm_nova
        perm_dist[k]<-perm_diff
    }
    #hist(perm_dist, main=z)
    num_ext<-length(which(perm_dist>q))
    p_val<-num_ext/N.PERM
    max_perm<-max(perm_dist)
    min_perm<-min(perm_dist)
    
    d<-list(line, min_perm, max_perm, q, num_ext, p_val)
    return(d)
}

outcome_E<-mapply(P_val_calc_E, E_clean.split, counts, Lines, line_obs_diff_E)
rownames(outcome_E) <- c("Line", "min_perm", "max_perm", "obs_diff", "per>obs", "p_val")
colnames(line_summary_data)<-Lines

write.table(outcome_E, "./E_permut.txt", sep="\t")


#summarize p-values and perform fdr correction
p_val_summary_E<-outcome_E[6,]
min(unlist(p_val_summary_E))
max(unlist(p_val_summary_E))
hist(unlist(p_val_summary_E))
p_E<-unlist(p_val_summary_E)

fdr_E <- p.adjust(p_E, method="fdr")
plot(line_obs_diff_E,p_E)
plot(line_obs_diff_E,fdr_E)
plot(p_E,fdr_E)

write.table(fdr_E, "./E_fdr.txt", sep="\t")

#conclusion: 175 (DN0748x37) looks like nova

################
#tan

library(readxl)
amer_genos <- read_excel("amer_genos.xlsx")

amer_genos$yellow <- factor(amer_genos$yellow)
amer_genos$tan <- factor(amer_genos$tan)
amer_genos$ebony <- factor(amer_genos$ebony)
amer_genos$Line <- factor(amer_genos$Line)

#divide datafile by gene and remove flies with missing genotypes
T<-subset(amer_genos, select=c(Line, Phenotype, tan))#pulls out data relevant for tan
T_clean<-na.omit(T) #removes flies with missing genotype data for tan
T_clean.split <- split(T_clean$Phenotype,T_clean$Line) #pulls out list of phenotypes for each line


#gets a list of Lines with tan genotypes, n=51 for tan
Lines<-unique(T_clean$Line)
Lines<-sort(Lines)
Lines<-as.character(Lines)


#summarize information about tan genotypes/phenotypes for each line
LINE_SUMMARY<- function(x) {
    line<-x
    obs_diff<-mean(T_clean[["Phenotype"]] [which(T_clean[["Line"]]==x & T_clean[["tan"]]=="0")]) - mean(T_clean[["Phenotype"]][which(T_clean[["Line"]]==x & T_clean[["tan"]]=="1")])
    n_amer<-NROW(T_clean[["Phenotype"]] [which(T_clean[["Line"]]==x & T_clean[["tan"]]=="0")])
    n_nova<-NROW(T_clean[["Phenotype"]][which(T_clean[["Line"]]==x & T_clean[["tan"]]=="1")])
    n_tot<-NROW(T_clean[["Phenotype"]][which(T_clean[["Line"]]==x)])
    sum_amer<-sum(T_clean[["Phenotype"]] [which(T_clean[["Line"]]==x & T_clean[["tan"]]=="0")])
    sum_nova<-sum(T_clean[["Phenotype"]][which(T_clean[["Line"]]==x & T_clean[["tan"]]=="1")])
    sum_tot<-sum(T_clean[["Phenotype"]][which(T_clean[["Line"]]==x)])
    avg_amer<-sum_amer/n_amer
    avg_nova<-sum_nova/n_nova
    avg_tot<-sum_tot/n_tot
    obs_diff<-avg_amer - avg_nova
    
    results<- list(line, n_amer, n_nova, n_tot, sum_amer, sum_nova, sum_tot, avg_amer, avg_nova, avg_tot, obs_diff)
    return(results)
}

line_summary_data<-sapply(Lines, LINE_SUMMARY, simplify=TRUE, USE.NAMES=FALSE)
rownames(line_summary_data) <- c("Line", "n_amer", "n_nova", "n_tot", "sum_amer", "sum_nova", "sum_tot", "avg_amer", "avg_nova", "avg_tot", "obs_diff")
colnames(line_summary_data)<-Lines
write.table(line_summary_data, "./T_line_summary_data.txt", sep="\t")

line_obs_diff_T<-line_summary_data[11,] #summary of observed differences by line
counts<-line_summary_data[2,] #number of D. americana alleles by line


#Define permutation values and pvalue collection list
p_val<- rep(NA, length(Lines))
N.PERM=10000

#resample function
P_val_calc_T<-function(x,y,z,q) #x=T_clean.split (pigmentation scores by line), y = number of americana genos per line, z = lines, q=obs_diff
{
    perm_dist <- rep(NA,N.PERM) #resest list for permuted values of difference between amer and nova
    line<-z
    for (k in 1:N.PERM)
    {
        sum_perm_amer<-sum(sample(x,y, replace=FALSE, prob=NULL)) #sample without replacement, equal frequency
        mean_perm_amer<-sum_perm_amer/y
        obs_sum<-sum(x) #sum of pig score for line
        sample_size<-length(x) #sample size for line
        sum_perm_nova<-obs_sum-sum_perm_amer
        mean_perm_nova<-sum_perm_nova/(sample_size-y)
        perm_diff<-mean_perm_amer - mean_perm_nova
        perm_dist[k]<-perm_diff
    }
    hist(perm_dist, main=z)
    num_ext<-length(which(perm_dist>q))
    p_val<-num_ext/N.PERM
    max_perm<-max(perm_dist)
    min_perm<-min(perm_dist)
    
    d<-list(line, min_perm, max_perm, q, num_ext, p_val)
    return(d)
}

outcome_T<-mapply(P_val_calc_T, T_clean.split, counts, Lines, line_obs_diff_T)
rownames(outcome_T) <- c("Line", "min_perm", "max_perm", "obs_diff", "per>obs", "p_val")
colnames(line_summary_data)<-Lines

write.table(outcome_T, "./T_permut.txt", sep="\t")


#summarize p-values and perform fdr correction
p_val_summary_T<-outcome_T[6,]
min(unlist(p_val_summary_T))
max(unlist(p_val_summary_T))
hist(unlist(p_val_summary_T))
p_T<-unlist(p_val_summary_T)

fdr_T <- p.adjust(p_T, method="fdr")
plot(line_obs_diff_T,p_T)
plot(line_obs_diff_T,fdr_T)
plot(p_T,fdr_T)

write.table(fdr_T, "./T_fdr.txt", sep="\t")

#conclusion: 118 (DA0626) looks like nova (P = 0.16)
# additional lines with marginal similarity to nova (P = 0.05 or P = 0.06): G9647, FP9918, II0710, SC0708, SC0718


################
#yellow

library(readxl)
amer_genos <- read_excel("amer_genos.xlsx")

amer_genos$yellow <- factor(amer_genos$yellow)
amer_genos$tan <- factor(amer_genos$tan)
amer_genos$ebony <- factor(amer_genos$ebony)
amer_genos$Line <- factor(amer_genos$Line)

#divide datafile by gene and remove flies with missing genotypes
Y<-subset(amer_genos, select=c(Line, Phenotype, yellow))#pulls out data relevant for yellow
Y_clean<-na.omit(Y) #removes flies with missing genotype data for yellow
Y_clean.split <- split(Y_clean$Phenotype,Y_clean$Line) #pulls out list of phenotypes for each line
Y_clean.split['33']<-NULL
Y_clean.split['36']<-NULL


#gets a list of Lines with yellow genotypes, n=49 for yellow
Lines<-unique(Y_clean$Line) 
Lines<-sort(Lines)
Lines<-as.character(Lines)


#summarize information about yellow genotypes/phenotypes for each line
LINE_SUMMARY<- function(x) {
	line<-x
	obs_diff<-mean(Y_clean[["Phenotype"]] [which(Y_clean[["Line"]]==x & Y_clean[["yellow"]]=="0")]) - mean(Y_clean[["Phenotype"]][which(Y_clean[["Line"]]==x & Y_clean[["yellow"]]=="1")])
	n_amer<-NROW(Y_clean[["Phenotype"]] [which(Y_clean[["Line"]]==x & Y_clean[["yellow"]]=="0")])
	n_nova<-NROW(Y_clean[["Phenotype"]][which(Y_clean[["Line"]]==x & Y_clean[["yellow"]]=="1")])
	n_tot<-NROW(Y_clean[["Phenotype"]][which(Y_clean[["Line"]]==x)])
	sum_amer<-sum(Y_clean[["Phenotype"]] [which(Y_clean[["Line"]]==x & Y_clean[["yellow"]]=="0")])
	sum_nova<-sum(Y_clean[["Phenotype"]][which(Y_clean[["Line"]]==x & Y_clean[["yellow"]]=="1")])
	sum_tot<-sum(Y_clean[["Phenotype"]][which(Y_clean[["Line"]]==x)])
	avg_amer<-sum_amer/n_amer
	avg_nova<-sum_nova/n_nova
	avg_tot<-sum_tot/n_tot
	obs_diff<-avg_amer - avg_nova
	
	results<- list(line, n_amer, n_nova, n_tot, sum_amer, sum_nova, sum_tot, avg_amer, avg_nova, avg_tot, obs_diff)
	return(results)
	}

line_summary_data<-sapply(Lines, LINE_SUMMARY, simplify=TRUE, USE.NAMES=FALSE)
rownames(line_summary_data) <- c("Line", "n_amer", "n_nova", "n_tot", "sum_amer", "sum_nova", "sum_tot", "avg_amer", "avg_nova", "avg_tot", "obs_diff")
colnames(line_summary_data)<-Lines
write.table(line_summary_data, "./Y_line_summary_data.txt", sep="\t")

line_obs_diff_Y<-line_summary_data[11,] #summary of observed differences by line
counts<-line_summary_data[2,] #number of D. americana alleles by line


#Define permutation values and pvalue collection list
p_val<- rep(NA, length(Lines))
N.PERM=10000

#resample function
P_val_calc_Y<-function(x,y,z,q) #x=Y_clean.split (pigmentation scores by line), y = number of americana genos per line, z = lines, q=obs_diff
	{ 
	perm_dist <- rep(NA,N.PERM) #resest list for permuted values of difference between amer and nova
	line<-z
for (k in 1:N.PERM) 
	{
	sum_perm_amer<-sum(sample(x,y, replace=FALSE, prob=NULL)) #sample without replacement, equal frequency
	mean_perm_amer<-sum_perm_amer/y
	obs_sum<-sum(x) #sum of pig score for line
	sample_size<-length(x) #sample size for line
	sum_perm_nova<-obs_sum-sum_perm_amer
	mean_perm_nova<-sum_perm_nova/(sample_size-y)
	perm_diff<-mean_perm_amer - mean_perm_nova
	perm_dist[k]<-perm_diff
} 
	hist(perm_dist, main=z)
	num_ext<-length(which(perm_dist>q))
	p_val<-num_ext/N.PERM
	max_perm<-max(perm_dist)
	min_perm<-min(perm_dist)

	d<-list(line, min_perm, max_perm, q, num_ext, p_val)
	return(d)
}	

outcome_Y<-mapply(P_val_calc_Y, Y_clean.split, counts, Lines, line_obs_diff_Y)
rownames(outcome_Y) <- c("Line", "min_perm", "max_perm", "obs_diff", "per>obs", "p_val")
colnames(line_summary_data)<-Lines

write.table(outcome_Y, "./Y_permut.txt", sep="\t")


#summarize p-values and perform fdr correction
p_val_summary_Y<-outcome_Y[6,]
min(unlist(p_val_summary_Y))
max(unlist(p_val_summary_Y))
hist(unlist(p_val_summary_Y))
p_Y<-unlist(p_val_summary_Y)

fdr_Y <- p.adjust(p_Y, method="fdr")
plot(line_obs_diff_Y,p_Y)
plot(line_obs_diff_Y,fdr_Y)
plot(p_Y,fdr_Y)

write.table(fdr_Y, "./Y_fdr.txt", sep="\t")

#conclusion: all are functionally equivalent to nova

##############################################
## [4] Test segregating sites for an association with pigmentation
## Variable sites with minor allele present in less than 5 individuals excluded.
## Sites with indels excluded.
## tan_sites_noindel.csv is provided as Supplementary Table 6.
## ebony_sites_noindel.csv is provided as Supplementary Table 7.
## Summary of output from general linear models (glms) for both genes provided in Supplementary Table 8.

library(seqinr)

tan_sites_noindel <- read_csv("tan_sites_noindel.csv",
col_types = cols(Pigmentation = col_number()))

ebony_sites_noindel <- read_csv("ebony_sites_noindel.csv",
col_types = cols(Pigmentation = col_number()))

# set col_number to chose columns to convert to factor
##change to factor
col_number<-c(3:42)
ebony_sites_noindel[,col_number]<-lapply(ebony_sites_noindel[,col_number],factor)

##some sites return "NA" because of genotypes perfectly correlated with another site.

tan_test<-glm(Pigmentation ~ site100+site1006+site1016+site1052+site1089+site1165+site1167+site13+site1300+site1301+site1302+site1303+site1316+site143+site155+site168+site176+site184+site189+site191+site205+site21+site216+site218+site228+site24+site247+site253+site267+site293+site296+site298+site305+site310+site311+site315+site319+site342+site344+site348+site350+site363+site365+site367+site379+site387+site403+site404+site412+site418+site422+site427+site441+site442+site472+site481+site516+site520+site522+site529+site543+site545+site551+site580+site615+site642+site658+site693+site720+site721+site736+site777+site825+site962, data=tan_sites_noindel)

summary(tan_test)

eb_test<-glm(Pigmentation ~ site120+site542+site457+site387+site477+site470+site468+site299+site318+site439+site447+site434+site450+site433+site432+site436+site458+site428+site463+site556+site497+site319+site452+site465+site83+site321+site386+site554+site557+site102+site317+site320+site366+site384+site34+site76+site129+site471+site68+site74, data=ebony_sites_noindel)

summary(eb_test)


################################
## [5] Correspondence analysis (CA) used to compare distributions of pigmentation
## phenotypes in backcross (BC1) populations among strains.
## CA plot produced with code below was color-coded manually in Adobe Illustrator to make Figure 4.

#convert data to univariate format
library(tidyr)
pig_classes.long<-pivot_longer(pig_classes,cols=3:10,names_to="Pigmentation_score",values_to = "Flies")
pig_classes.univ<-uncount(pig_classes.long, Flies)

#Correspondence analysis (CA)
library("FactoMineR")
library("factoextra")

#data: strain, number of flies in categories 1-8
res.ca <- CA(pig_classes.str.df, quali.sup=1, graph = FALSE)
fviz_ca_biplot(res.ca, repel = TRUE)

#get data from CA for strains
strains_ca<-get_ca_row(res.ca)

#pull out coordinates table
ca_coord<-strains_ca$coord

#append to original datafile to get strain names
coord<-cbind(pig_classes.str.df,ca_coord)

#plot CA coordinates, labeled by strain
ggplot(coord,aes(x=`Dim 1`,y=`Dim 2`))+geom_point()+geom_text(aes(label=`Strain`),hjust=0,vjust=0)

#calculate Euclidian distance in Dim 1/Dim 2 space between all pairs of strains
library(distances)

#prepare data frame
dist_coord<-subset(coord, select=c(1,10,11))

#calculate all pairwise Euclidean distances
my_distances2 <- distances(dist_coord, id_variable = "Strain")

#unpack to univariate format
library(reshape2)
results_dist_coord <- melt(as.matrix(my_distances2), varnames = c("row", "col"))

#remove comparisons to self
results_dist_coord<-results_dist_coord[!(results_dist_coord$value==0),]

#export Distances summary
write.csv(results_dist_coord, "Distances.txt")

#add population column indicating whether each pair is from the same or different collection site in Excel.
#(yes, I should learn to use R better...) and reimport modified file
 Distances <- read_csv("Distances.csv")

#inspect distributions of distances for pairs from same and different collection sites - look very similar
ggplot(Distances,aes(x=value,group=population,fill=population))+
+     geom_histogram(position="dodge",binwidth=0.25)+theme_bw()

#make subsets containing distances only for comparisons between strains from "same" and "different" populations
same<-subset(Distances, population == 'same',)
different<-subset(Distances, population == 'different',)

#compare two distributions with t.test
t.test(same$value,different$value)

#output: Welch Two Sample t-test
# t = 0.7647, df = 118.82, p-value = 0.446
# alternative hypothesis: true difference in means is not equal to 0
# 95 percent confidence interval: -0.04209808  0.09507055
# sample estimates:mean of x mean of y
# 0.6808331 0.6543469
